# Continental-scale multi-omics reveals a distinct microbial-viral biome in sandy beach ecosystems structured by tidal zonation

**DOI:** 10.64898/2026.04.10.717577

**Authors:** Xiyang Dong, Feng Cai, Yingchun Han, Chuwen Zhang, Hongshuai Qi, Shaohua Zhao, Liang Wang, Zhong Pan, Yanbin Chen, Zhengran Li, Zijian Lu, Xiaomei Guo, Yiting Ji, Jianhui Liu, Shengjie Li, Chujin Ruan, Lu Zhang

**Affiliations:** Third Institute of Oceanography, Ministry of Natural Resources, Xiamen 361005, China; College of Ocean & Earth Sciences, Xiamen University, Xiamen 361102, China; State Key Laboratory of Submarine Geoscience, and School of Oceanography, Shanghai Jiao Tong University, Shanghai 200240, China; Department of Health and Environmental Sciences, School of Science, Xi’an Jiaotong-Liverpool University, Suzhou 215123, China

## Abstract

Sandy beaches are dynamic coastal interfaces shaped by strong physical forcing and intense exchange between marine and terrestrial environments, yet their microbiomes remain poorly resolved at the genomic scale. Here we present a genome-resolved survey of microbial and viral communities across sandy beaches spanning a continental-scale latitudinal gradient along the Chinese coastline. By integrating cross-shore sampling, coastal geochemistry and large-scale multi-omics, we generated 978 metagenomes, 63 viromes and 72 metatranscriptomes, reconstructing 13,337 metagenome-assembled genomes and 38,255 viral populations. Sandy beach microbiomes exhibit exceptionally high genomic novelty, with more than 90% of species-level genomes representing previously undescribed taxa, suggesting that permeable coastal sediments constitute a distinct microbial and viral reservoir. Tidal zonation emerged as a dominant ecological driver structuring microbial diversity, metabolic strategies and virus-host interactions across cross-shore gradients. Genome-resolved analyses revealed systematic metabolic shifts from oxic heterotrophy in supratidal sediments toward increasingly chemolithotrophic and autotrophic pathways toward the low-intertidal and subtidal zone. Sandy beach microbiomes further encode broad potential for hydrocarbon and plastic transformation, together with diverse biosynthetic and antibiotic resistance repertoires that may mediate microbial chemical interactions. Together, these findings identify sandy beaches as a previously under-recognized microbial-viral biome shaped by tidal forcing, providing insight into microbiome evolution and coastal ecosystem resilience under increasing anthropogenic pressure.

## Introduction

Sandy beaches occupy approximately one-third of the world’s ice-free coastline and represent one of the most intensively used land-sea interfaces^1^. Despite their apparent geomorphic simplicity, sandy beaches are highly dynamic systems shaped by waves, tides, and sediment transport that continually reorganize pore structure and hydrological connectivity across spatial and temporal scales. Permeable sands can function as advective biogeochemical reactors, facilitating rapid exchange among seawater, groundwater and the atmosphere^2^. Tidal pumping and wave-driven circulation generate steep gradients in oxygen availability, redox potential and nutrient concentrations, enabling intense microbial processing of carbon, nitrogen and trace elements before solutes are returned to coastal waters^3–5^. Because tidal elevation determines the frequency and duration of inundation, desiccation and oxygen exposure, cross-shore zonation provides a predictable physical template that may structure microbial metabolism and biogeochemical transformations along the land-sea interface.

Despite the recognized importance of permeable sediments for coastal biogeochemistry, the microbial and viral communities inhabiting sandy beaches remain poorly resolved, particularly at the genomic scale^6–8^. Previous studies have shown that microbial assemblages vary across intertidal microhabitats, reflecting gradients in hydrodynamic forcing, redox conditions and resource availability^9–14^. However, most investigations rely on single-site surveys or marker-gene approaches, limiting understanding of whether consistent ecological patterns emerge across coastlines spanning different climatic regimes, morphodynamic settings and environmental pressures. This limitation reflects the intrinsic properties of permeable beach sands, where rapid wetting-drying cycles, salinity fluctuations and ultraviolet exposure coincide with low biomass, strong microscale heterogeneity and advective redox oscillations, jointly selecting for metabolically flexible microorganisms while complicating genome-resolved characterization^15^. Viruses influence microbial mortality, horizontal gene transfer and host metabolic potential in many ecosystems^16^. However, they remain especially understudied in permeable coastal sediments, leaving unresolved how virus-host interactions contribute to microbial adaptation and biogeochemical functioning across tidal gradients.

Human activity introduces an additional layer of environmental complexity. Sandy beaches are among the most heavily visited coastal ecosystems and receive diverse anthropogenic inputs, including nutrients, trace metals, antibiotics, hydrocarbons and microplastics^16, 17^. These compounds may alter microbial resource availability, impose selective pressures on resistance and detoxification mechanisms, and influence horizontal gene transfer^18, 19^. Because contaminant accumulation and transport depend strongly on hydrodynamic flushing, sediment permeability and residence time, anthropogenic impacts are likely modulated by the same physical processes that structure microbial habitats. Understanding how physical forcing, environmental chemistry and human pressures interact to shape sandy-beach microbiomes therefore represents an important challenge for predicting coastal ecosystem functioning and resilience.

Here we present a continental-scale genome-resolved investigation of sandy beach microbiomes and viromes across 18 beaches spanning tropical to temperate climatic zones, contrasting morphodynamic settings and varying levels of human use along the Chinese coastline. By integrating cross-shore sampling, coastal chemistry and large-scale multi-omics analyses spanning sediment and water samples, we generated 978 metagenomes, 63 viromes and 72 metatranscriptomes, enabling reconstruction of 13,337 metagenome-assembled genomes and 38,255 viral populations. Our analyses identify sandy beaches as a distinct microbial-viral biome characterized by high genomic novelty, strong tidal structuring and broad functional potential for organic matter turnover, redox flexibility and pollutant transformation. These results provide a genomic framework for understanding how physical forcing structures microbiomes in permeable coastal sediments and establish a reference resource for predicting ecosystem responses to anthropogenic pressure and climate-driven coastal change.

## Results

### Environmental gradients across continental-scale sandy beach ecosystems

To characterize environmental variation across sandy beach ecosystems, we conducted a coordinated survey along the Chinese coastline spanning a latitudinal gradient from 18°N to 40°N. In total, 18 sandy beaches were investigated, including 15 urban recreational beaches located in major coastal tourism cities and three natural beaches serving as reference sites (**Fig. 1a–b and Tables S1–S2**). These sites span tropical, subtropical and temperate climatic zones along the South China Sea, East China Sea, Yellow Sea and Bohai Sea coasts, capturing broad environmental variation across the Chinese coastline.

**Figure 1.**
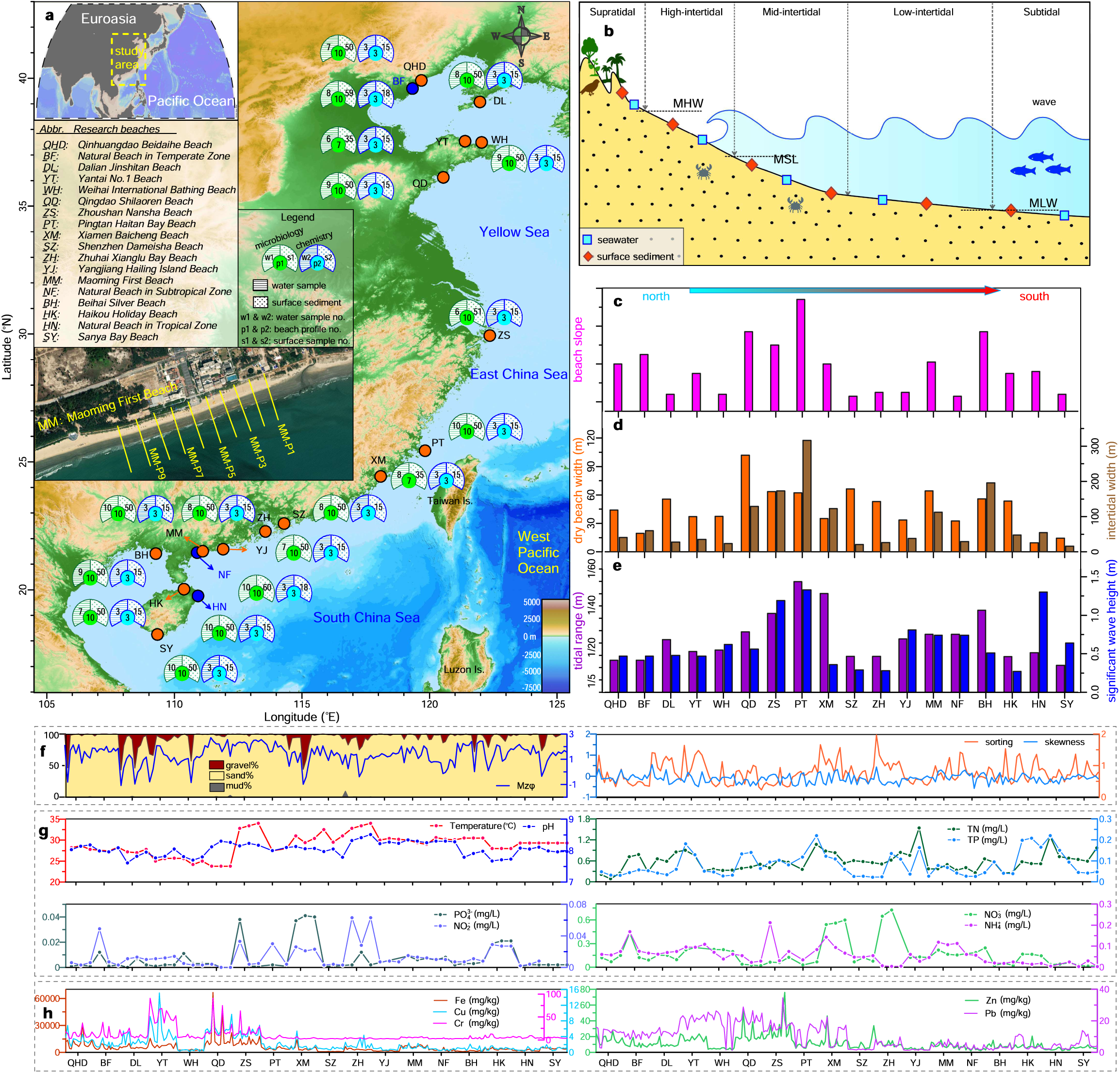
Study sites, sampling design and environmental gradients across sandy beach ecosystems along the Chinese coastline. **(a)** Locations of 18 sandy beaches (18°–40°N) spanning tropical, subtropical and temperate zones, including 15 recreational beaches and three natural reference sites. Circular charts indicate the numbers of water and sediment samples collected at each site. **(b)** Conceptual cross-shore sampling framework showing the supratidal, high-intertidal, mid-intertidal, low-intertidal and subtidal zones, together with the positions of surface sediment and seawater sampling. MHW, mean high water; MSL, mean sea level; MLW, mean low water. **(c)** Beachface slope variation among sites. **(d)** Cross-shore geomorphological characteristics across beaches, including dry-beach width (orange bars) and intertidal width (brown bars). **(e)** Tidal range (purple bars) and significant wave height (blue bars) across study sites. **(f**) Surface sediment grain-size composition (gravel, sand and mud fractions) and sediment textural parameters, including mean grain size, sorting and skewness. **(g)** Water physicochemical variables, including temperature, pH and dissolved nutrients. **(h)** Metal concentrations (Fe, Cu, Zn, Pb and Cr) in sediments across sites.

The surveyed beaches displayed pronounced variation in geomorphology and hydrodynamic regimes (**Fig. 1c–f and Tables S3–S4**). Based on beachface slope and sediment characteristics, the beaches were classified into reflective, intermediate and dissipative morphodynamic types. Beachface slopes ranged from steep reflective systems (∼1/8–1/10) to gently sloping dissipative beaches (<1/40) (**Fig. 1c**). Dry-beach widths varied from ∼14 to 102 m, whereas intertidal widths ranged from ∼15 m to more than 300 m, indicating substantial cross-site variation in beach morphology (**Fig. 1d**). Hydrodynamic forcing also varied among sites, with tidal ranges of approximately 1.0–4.5 m and significant wave heights of ∼0.28–1.33 m (**Fig. 1e**). Surface sediments were dominated by sand fractions across all sites, typically exceeding 80% of total sediment composition, with minor contributions from gravel and mud (**Fig. 1f**).

Environmental conditions varied markedly across the coastline. Surface-water temperature ranged from 23.8 °C in northern temperate regions to 34.0 °C at southern tropical beaches, whereas pH remained consistently alkaline (7.67–8.51) (**Fig. 1g and Table S5**). Nutrient concentrations, including total nitrogen, total phosphorus and dissolved inorganic nitrogen and phosphorus species, also varied substantially among beaches. Metal concentrations (Fe, Cu, Zn, Pb and Cr) also showed strong spatial variability and clear partitioning between dissolved and sediment-associated fractions (**Fig. 1h and Tables S5-S6**). Metal concentrations were generally higher in sediments than in the overlying water column. Antibiotics were widely detected across beaches **(Table S7**). Concentrations in seawater typically occurred at ng L^-1^ levels, whereas sediment concentrations were generally observed at pg g^-1^ levels. Within individual beaches, antibiotic concentrations were often higher in mid-intertidal sediments than in upper beach zones, consistent with an influence of tidal exchange and hydrodynamic processes on contaminant accumulation in sandy beach sediments.

### Tidal zonation structures prokaryotic and viral communities across sandy beaches

To characterize community organization across sandy beach ecosystems, we analyzed 978 metagenomes together with 63 viromes, enabling integrated comparisons of prokaryotic and viral communities across tidal zones and environmental compartments. Across all samples, prokaryotic communities spanned 81 phyla, including 74 bacterial and 7 archaeal phyla (**Fig. 2a and Table S8**). Community composition was consistently dominated by *Pseudomonadota*, *Bacteroidota*, *Actinomycetota* and *Bacillota*, which together accounted for approximately 80% of total relative abundance across sites. Clear cross-shore patterns were observed: *Pseudomonadota* increased toward intertidal sediments, whereas *Actinomycetota* and *Bacillota* were relatively enriched in supratidal sands. Archaeal lineages, primarily *Thermoproteota* and *Thermoplasmatota*, were more abundant in lower intertidal sediments, consistent with stronger redox gradients and organic matter accumulation in periodically inundated zones^20^. Viral communities were dominated by fewer major taxonomic groups but were strongly dominated by *Caudoviricetes*, with additional contributions from *Faserviricetes*, *Megaviricetes* and *Tectiliviricetes* (**Fig. 2b, Fig. S1 and Table S9**). Viral assemblages also exhibited zonation across tidal habitats. *Faserviricetes* were more abundant in supratidal sands, whereas *Megaviricetes* contributed more prominently in several intertidal samples, although *Caudoviricetes* remained dominant across most sites.

**Figure 2.**
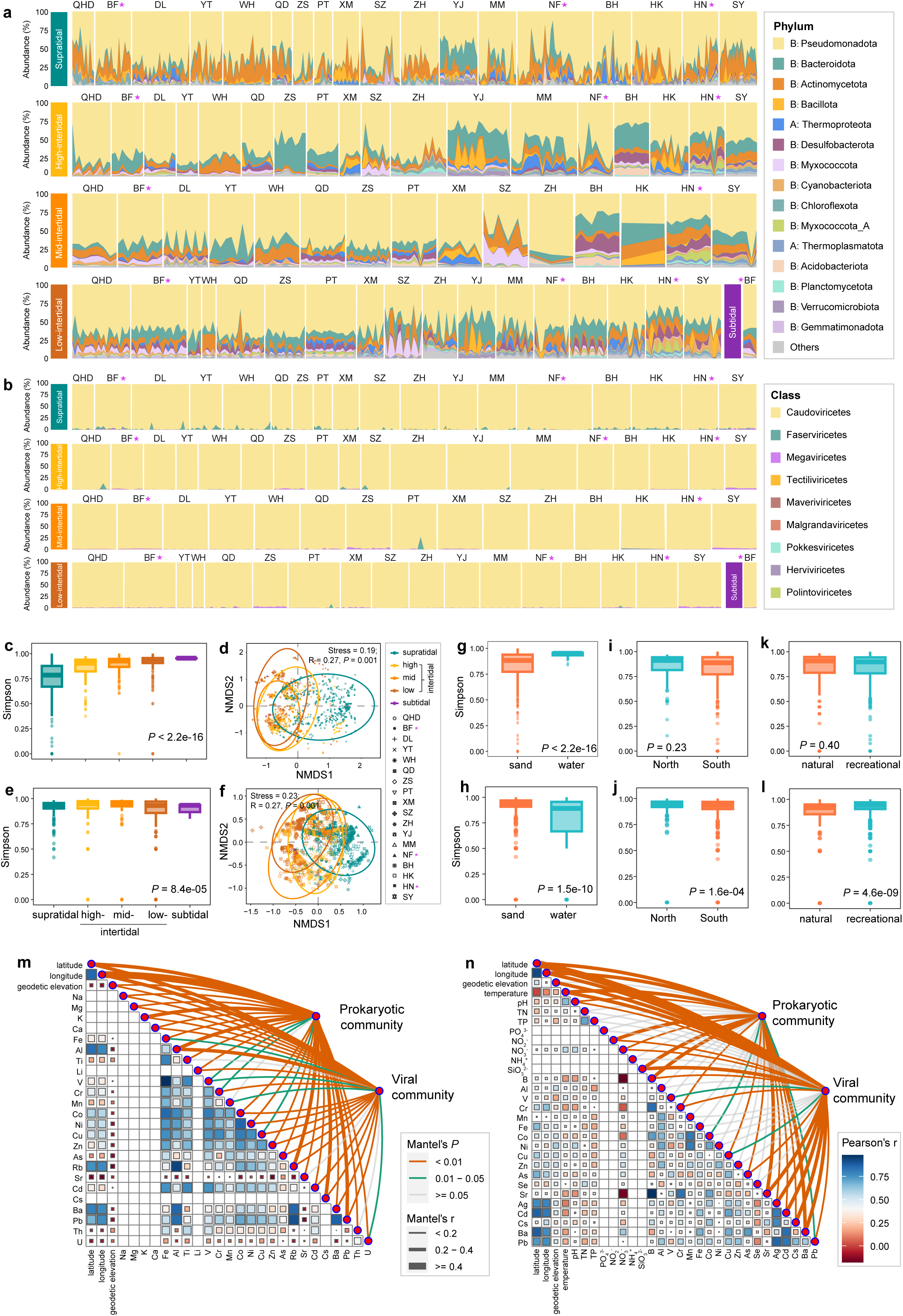
Composition, diversity and environmental drivers of prokaryotic and viral communities across sandy beaches. **(a, b)** Taxonomic composition of prokaryotic (phylum level) **(a)** and viral (class level) **(b**) communities across beaches, with samples grouped by tidal zone within each site (supratidal, high-intertidal, mid-intertidal, low-intertidal and subtidal). Sites are ordered latitudinally from north to south along the Chinese coastline. Asterisks denote natural reference beaches. **(c, d)** Prokaryotic α-diversity (Simpson index) across tidal zones **(c)** and β-diversity visualized by non-metric multidimensional scaling (NMDS) based on Bray–Curtis dissimilarities **(d)**. **(e, f)** Viral α-diversity (Simpson index) across tidal zones **(e)** and β-diversity visualized by NMDS based on Bray–Curtis dissimilarities **(f)**. **(g–l)** Comparisons of α-diversity among environmental groupings, including matrix type (sand versus water), geographic region (north versus south), and beach usage (natural versus recreational), for prokaryotic **(g, i, k)** and viral **(h, j, l)** communities. **(m, n)** Relationships between environmental variables and prokaryotic and viral community composition for sediment-associated geochemical variables **(m)** and water physicochemical variables together with selected elemental variables **(n)**. Heatmaps show Pearson correlations among environmental variables, and links indicate significant Mantel correlations between environmental factors and community composition. For boxplots, center lines indicate medians, boxes indicate interquartile ranges, and whiskers indicate 1.5 × interquartile ranges. α-diversity differences were assessed using Kruskal–Wallis tests or two-sided Wilcoxon rank-sum tests, as appropriate. β-diversity differences were evaluated using ANOSIM with 999 permutations. Exact statistics are provided in **Tables S10–S15**.

Across the coastline, tidal zonation emerged as the strongest driver of microbial community structure. Prokaryotic α-diversity increased progressively from supratidal to subtidal zones (Simpson index, *P* < 2.2 × 10^-16^; **Fig. 2c, Fig. S2 and Tables S10–S11**), consistent with reduced desiccation stress and enhanced hydrological exchange in frequently inundated sediments^11^. Correspondingly, β-diversity analyses revealed clear compositional differentiation among tidal zones (ANOSIM *R* = 0.27, *P* = 0.001; **Fig. 2d and Fig. S3a**), with the largest dissimilarities observed between supratidal and lower intertidal communities. Viral diversity showed a similar but not identical cross-shore pattern. Viral α-diversity peaked in mid- and low-intertidal sediments (**Fig. 2e, Fig. S4 and Tables S12–S13**), and viral community turnover closely paralleled prokaryotic compositional shifts across tidal zones (ANOSIM *R* = 0.27, *P* = 0.001; **Fig. 2f and Fig. S5a**). Habitat matrix further structured microbial communities. Prokaryotic α-diversity was higher in water samples than in sediments, whereas water samples showed a narrower range of Simpson diversity values (*P* < 2.2 × 10^-16^; **Fig. 2g and Fig. S3b**). In contrast, viral diversity was significantly higher in sediments than in water (*P* = 1.5 × 10^-10^; **Fig. 2h and Fig. S5b**), consistent with the possibility that porous sediments act as viral retention zones that enhance virus-host interactions^21^.

Geographic gradients imposed additional structure across the coastline. Although prokaryotic α-diversity did not differ significantly between northern and southern beaches (Simpson index, *P* = 0.23; **Fig. 2i**), community composition exhibited clear regional separation (ANOSIM *R* = 0.27, *P* = 0.001; **Fig. S3c**), suggesting distance-decay relationships and climatic filtering^9^. Viral communities showed stronger geographic differentiation, with higher α-diversity and greater compositional turnover observed in northern sites (**Fig. 2j and Fig. S5c**). In contrast, anthropogenic disturbance exerted comparatively modest effects. Prokaryotic communities did not show significant shifts in diversity or composition between natural and recreational beaches (**Fig. 2k and Fig. S3e–f**), suggesting limited detectable effects of beach usage on prokaryotic community diversity and composition in this dataset. Viral richness, however, was lower at natural reference sites (*P* = 4.6 × 10^-9^; **Fig. 2l and Fig. S5e–f**), indicating that beach usage may be associated with differences in local virus-host dynamics^13, 22^. Mantel tests revealed that both prokaryotic and viral community structures were significantly associated with environmental variables (**Fig. 2m–n and Tables S14–S15**). Geographic position, physicochemical parameters, nutrients and trace metals collectively contributed to microbial biogeographic patterns, with viral communities showing particularly strong correlations with nutrient and metal gradients^23^.

### Sandy beaches harbor extensive and largely distinct microbial genomic diversity

To characterize the genomic diversity of sandy beach microbiomes, we reconstructed microbial genomes from 978 metagenomic assemblies, generating a comprehensive genome catalogue spanning sandy beach sediments and adjacent water samples. After dereplication at 95% average nucleotide identity (ANI) and quality filtering according to MIMAG standards^24^, a total of 13,337 medium- to high-quality metagenome-assembled genomes (MAGs) were retained to construct the Sandbeach Microbiome Genome Catalogue (SMGC) (**Fig. 3 and Table S16**). These genomes exhibited a mean completeness of 86.7% and mean contamination of 2.4%, including 5,607 near-complete genomes, 6,494 high-quality genomes and 1,236 medium-quality genomes (**Fig. S6a**). Genome sizes ranged from 0.25 to 14.8 Mb, reflecting substantial life-history diversity among sandy beach microorganisms (**Fig. S6b**). The smallest genomes were primarily affiliated with ultrasmall bacterial lineages such as *Patescibacteriota*, whereas the largest genomes belonged to *Myxococcota*, a group known for complex predatory and social behaviors^25, 26^.

**Figure 3.**
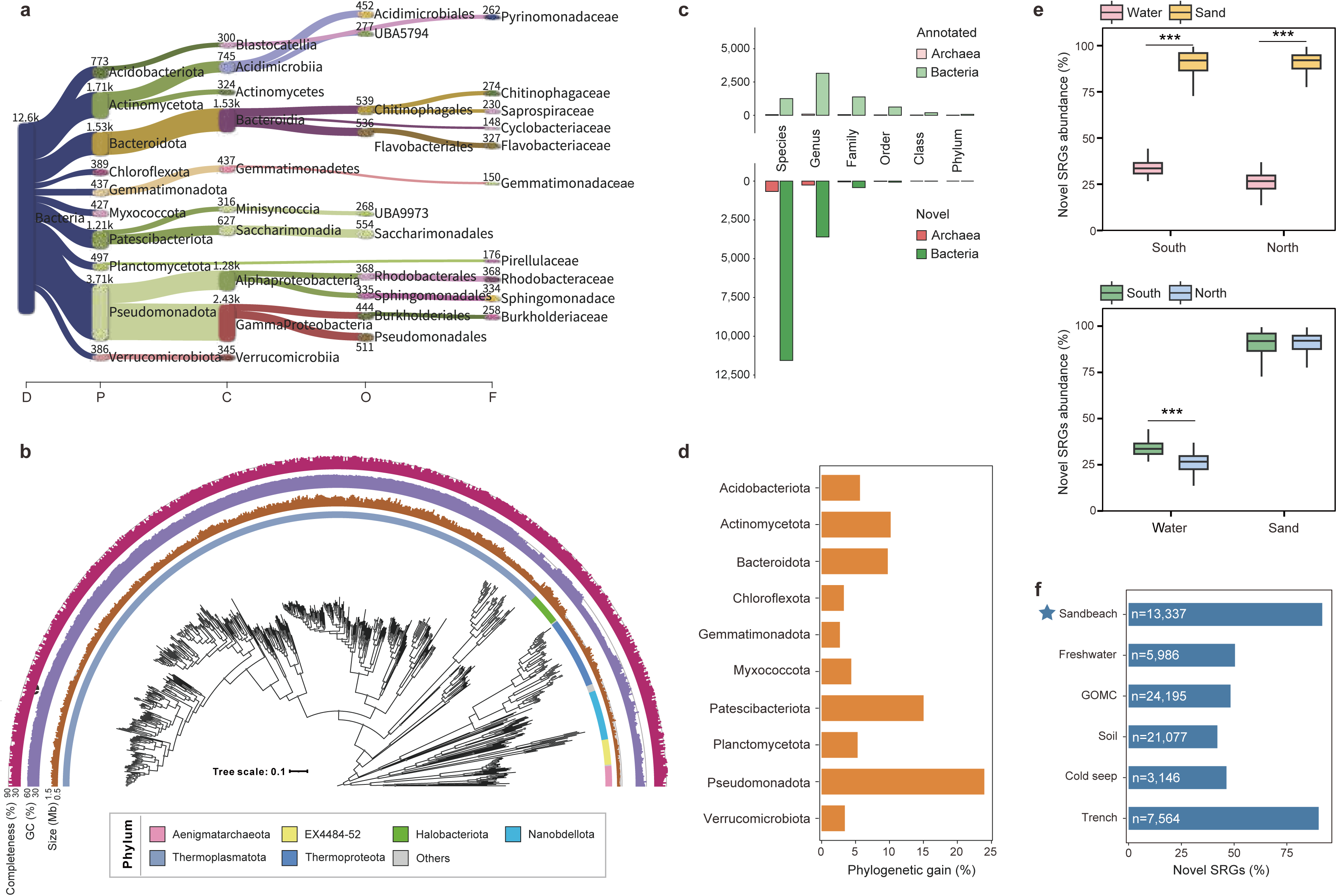
Taxonomic composition and genomic novelty of the Sandbeach Microbiome Genome Catalogue (SMGC). **(a)** Taxonomic distribution of bacterial metagenome-assembled genomes (MAGs) across taxonomic ranks from phylum to family based on GTDB r226. **(b)** Maximum-likelihood phylogenomic tree of archaeal MAGs reconstructed from 120 concatenated single-copy marker genes. Outer rings indicate phylum affiliation, genome size, GC content and genome completeness. **(c)** Taxonomic novelty relative to GTDB r226, showing the numbers of annotated and novel bacterial and archaeal lineages at each taxonomic rank. **(d)** Contribution of SMGC genomes to phylogenetic diversity within the ten most abundant bacterial phyla. Values indicate the percentage increase in phylogenetic diversity relative to the corresponding reference genome set. **(e)** Relative abundance of novel species-representative genomes (SRGs) across habitat types (water versus sand) and geographic regions (south versus north), shown separately for habitat and geographic comparisons. Boxplots show medians (center lines), interquartile ranges (boxes), and 1.5 × interquartile-range whiskers. Statistical significance was assessed using two-sided Wilcoxon rank-sum tests. **(f)** Comparison of species-level novelty between the SMGC and genome catalogues from Sandbeach, Freshwater, GOMC, Soil, Cold seep and Trench ecosystems. The number of genomes in each catalogue is indicated in white.

Taxonomic classification using the Genome Taxonomy Database (GTDB r226) revealed extensive phylogenetic diversity spanning 64 bacterial and 11 archaeal phyla (**Table S17**). Dominant bacterial phyla included *Pseudomonadota*, *Actinomycetota*, *Bacteroidota*, and *Patescibacteriota* (**Fig. 3a and Fig. S7**). Archaeal diversity was also substantial, with MAGs assigned to *Thermoproteota*, *Nanoarchaeota* and several additional archaeal lineages (**Figs. 3b and Fig. S8**). Despite this broad taxonomic representation, most genomes corresponded to previously undescribed species. Across major phyla, more than 80% of MAGs could not be assigned to known species-level taxa in the GTDB database (**Fig. 3c and Fig. S9**). Several dominant phyla, including *Pseudomonadota*, *Patescibacteriota*, *Actinomycetota* and *Bacteroidota*, showed substantial phylogenetic gains, ranging from 2.69% to 23.98% (**Fig. 3d**). Genomic novelty was strongly associated with habitat type. The relative abundance of novel species-representative genomes was significantly higher in sand-associated communities (mean 88.9%) than in overlying seawater (30.8%; *P* < 0.001) (**Fig. 3e**), suggesting that the heterogeneous microhabitats within sandy sediments may provide ecological niches that favor microbial diversification. Across tidal zones, novel species were most abundant in mid-intertidal sediments, where periodic tidal flushing and resource renewal create dynamic environmental conditions (**Fig. S10a**). In contrast, northern and southern regions showed no significant differences in novelty (**Fig. S10b**), suggesting that microhabitat heterogeneity within beaches exerts a stronger influence on microbial diversification than broad geographic gradients^10^.

To place sandy beach microbiomes in a broader ecological context, we compared the SMGC with genome catalogues from other major environments under a unified taxonomic framework (**Fig. 3f**). The SMGC exhibited an exceptionally high proportion of species-representative genomes representing previously uncharacterized taxa (91.7%), exceeding levels reported for freshwater^27^, soil^28^, and open-ocean microbiomes^29^ and approaching those observed in extreme deep-sea environments such as hadal trenches^30^. Consistent with this pattern, cross-habitat comparisons revealed remarkably limited genomic overlap between sandy beach microbiomes and other ecosystems. At a 95% ANI threshold, only 4.75% of SMGC genomes overlapped with oceanic datasets, whereas overlaps with soil and freshwater microbiomes were negligible (<0.25%; **Fig. S11a**). Most shared genomes originated from epipelagic seawater communities (**Fig. S11b**), consistent with microbial exchange between coastal waters and beach sediments driven by waves and tidal processes^31^. Together, these results indicate that sandy beaches harbor extensive and largely distinct microbial genomic diversity.

### Sandy beaches are a vast and previously unrecognized viral diversity hotspot

To characterize viral diversity across sandy beach ecosystems, we reconstructed viral genomes from 978 metagenomes together with 63 viromes (1,041 datasets in total), generating a large-scale viral genome collection from sandy beach sediments and adjacent coastal waters. In total, 207,056 viral contigs were identified and clustered into 38,255 viral operational taxonomic units (vOTUs) at 95% ANI, forming the Sandbeach Virome Catalogue (SVC) (**Fig. 4a, Fig. S12 and Table S18**). Genome completeness assessment indicated that the catalogue contains 6,627 complete, 6,683 high-quality, and 24,920 medium-quality viral genomes (**Fig. S13**). The majority of viral genomes ranged between 20 and 70 kb, consistent with genome sizes commonly reported for double-stranded DNA viruses infecting prokaryotes^32^.

**Figure 4.**
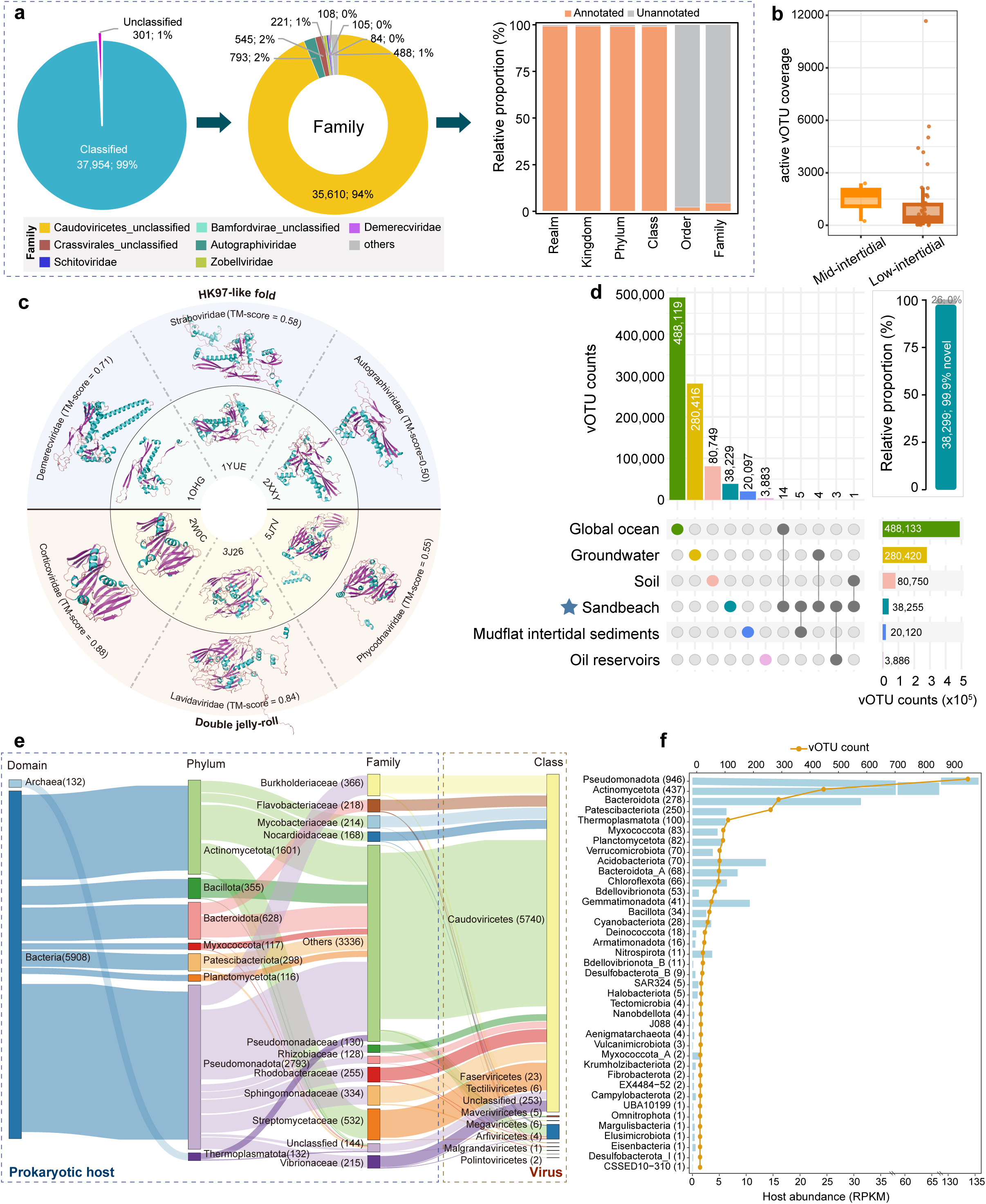
Taxonomic composition, activity, structural diversity and host associations of viruses in the Sandbeach Virome Catalogue (SVC). **(a)** Taxonomic annotation of viral operational taxonomic units (vOTUs) in the SVC. The left panel shows the proportions of classified and unclassified vOTUs. The middle panel displays the family-level composition of classified vOTUs, and the right panel summarizes the relative proportions of annotated and unannotated vOTUs across major taxonomic ranks from realm to family. **(b)** Transcript-derived coverage of active vOTUs in water samples from the mid- and low-intertidal zones. **(c)** Structural diversity of major capsid proteins (MCPs). Predicted MCP models were aligned to representative reference structures, revealing double jelly-roll and HK97-like folds. **(d)** Overlap between the SVC and virome catalogues from other environments, showing shared and habitat-specific vOTUs together with total catalogue sizes. Similarity thresholds used for cross-catalogue comparisons are described in the Methods. (**e)** Sankey diagram summarizing predicted virus–host linkages from prokaryotic domain, phylum and family to viral class. **(f)** Comparison between host phylum abundance and the number of linked vOTUs. Blue bars indicate mean host abundance (RPKM) across sandy beach samples, and the orange line indicates the number of vOTUs predicted to infect each host phylum. Detailed taxonomic, structural and virus–host association data are provided in **Tables S19–S23**.

Taxonomic annotation revealed that the vast majority of vOTUs belonged to the viral realm *Duplodnaviria*, with the class *Caudoviricetes* overwhelmingly dominating the viral assemblage (**Fig. 4a, Fig. S14 and Table S19**). Despite this dominance, taxonomic resolution beyond the class level remained extremely limited. More than 95% of annotated vOTUs could not be assigned to currently recognized viral families, underscoring the extensive unexplored diversity of viruses inhabiting sandy beach sediments. Metatranscriptomic analyses revealed transcriptional activity for a subset of vOTUs in intertidal water samples. In total, 2,255 vOTUs exhibited detectable transcriptional activity across the dataset (**Table S20**). Transcriptionally active viruses were detected across both tidal zones, with higher transcriptional coverage observed in water samples from the mid-intertidal zone (**Fig. 4b and Fig. S15**), suggesting enhanced virus-host interactions in frequently inundated regions of the beach.

Structural annotation of viral proteins further highlighted extensive diversity in capsid architectures (**Fig. 4c and Table S21**). A total of 367,320 major capsid proteins (MCPs) were identified across the SVC, most of which displayed the canonical HK97-like fold characteristic of tailed bacteriophages (n = 313,093). A smaller subset of MCPs exhibited double jelly-roll (DJR) folds (n = 2,657), indicating structural diversity and potential evolutionary links among distinct viral lineages. The remaining MCPs could not be confidently assigned to known structural folds, suggesting the presence of additional, yet uncharacterized capsid architectures^33^. To assess habitat specificity, the SVC was compared with viral genome catalogues from multiple ecosystems, including global ocean viromes^34^, groundwater^35^, soils^36^, mudflat sediments^23^ and oil reservoirs^37^ (**Fig. 4d**). Remarkably, only 26 vOTUs were shared across all environments, and overlaps between sandy beach viruses and any individual ecosystem were extremely limited. These patterns suggest that sandy beach viral communities represent a largely distinct viral reservoir shaped by the unique physicochemical conditions and hydrodynamic disturbances characteristic of permeable coastal sediments^38^. Viral replication strategies were predicted for 6,540 vOTUs using deep-learning approaches^39^ (**Fig. S14c and Table S22**). Among these, lytic viruses (4,561 vOTUs) were more prevalent than temperate viruses (1,979 vOTUs), consistent with the possibility that rapid host turnover in dynamic sandy beach environments favors lytic infection strategies^40^.

Integration of viral genomes with microbial MAGs enabled reconstruction of virus-host interaction networks. A total of 6,858 virus-host linkages were identified, connecting 6,655 vOTUs to 4,472 microbial genomes (**Fig. 4e and Table S23**). Predicted hosts spanned 41 bacterial and 9 archaeal phyla, with most linkages associated with dominant bacterial groups including *Pseudomonadota*, *Actinomycetota*, and *Bacteroidota*. Notably, the distribution of virus-host associations did not scale directly with host abundance across phyla (**Fig. 4f**), indicating that viral host range and infection patterns are shaped by ecological interactions beyond simple host availability^41^. Furthermore, based on CRISPR-informed iPHoP predictions^42^, 14.6% of virus-host clusters contained more than one viral or host genome, suggesting that multi-host associations and shared host ranges occur in sandy beach viral networks^43^.

### Tidal forcing drives metabolic stratification across sandy beach habitats

To characterize the metabolic potential of sandy beach microbiomes, we constructed a nonredundant gene catalogue comprising more than 4.08 × 10^8^ protein-coding gene clusters, a gene-catalogue size exceeding those reported for several previously published oceans^44–46^, soils^47^, and glacier microbiome datasets^48^ (**Fig. S16a**). Despite annotation with KEGG, eggNOG, Pfam, and CAZy, approximately 40% of gene clusters remained functionally uncharacterized (unannotated and of unknown function; **Fig. S16**), indicating substantial unexplored metabolic potential^45^. Functional profiles were dominated by heterotrophic carbon utilization and energy metabolism, with genes related to carbohydrate and amino acid metabolism, energy production, and inorganic ion transport accounting for >60% of annotated functions (**Fig. S16c**).

Genome-resolved analyses revealed a clear reorganization of microbial metabolic strategies along the tidal gradient, with oxic heterotrophic processes dominating upper beach sediments and chemolithotrophic and autotrophic pathways becoming increasingly important toward the subtidal zone (**Fig. 5a and Table S24**). Genes associated with complex organic matter degradation, including starch (*amyA*; mean 102 GPM), hemicellulose (*manA*; mean 99 GPM) and cellulose degradation (*bglB*; mean 106 GPM), were widely distributed and generally showed higher abundances in supratidal and upper intertidal sediments (**Fig. S17a**). These substrates are further processed through central carbon metabolism, as reflected by the widespread presence of glycolysis (*pfkA*; mean 99 GPM) and tricarboxylic acid cycle genes (*gltA*; mean 417 GPM). Together with the high abundance of aerobic respiration genes (**Fig. S17b**), particularly cytochrome c oxidase (*coxA*; mean 584 GPM), these patterns indicate that oxic heterotrophic metabolism represents the primary energy-generating strategy in upper beach sediments. These pathways were primarily associated with bacterial phyla including *Pseudomonadota*, *Bacteroidota* and *Actinomycetota*, which together contributed the majority of genes involved in organic carbon degradation and aerobic respiration.

**Figure 5.**
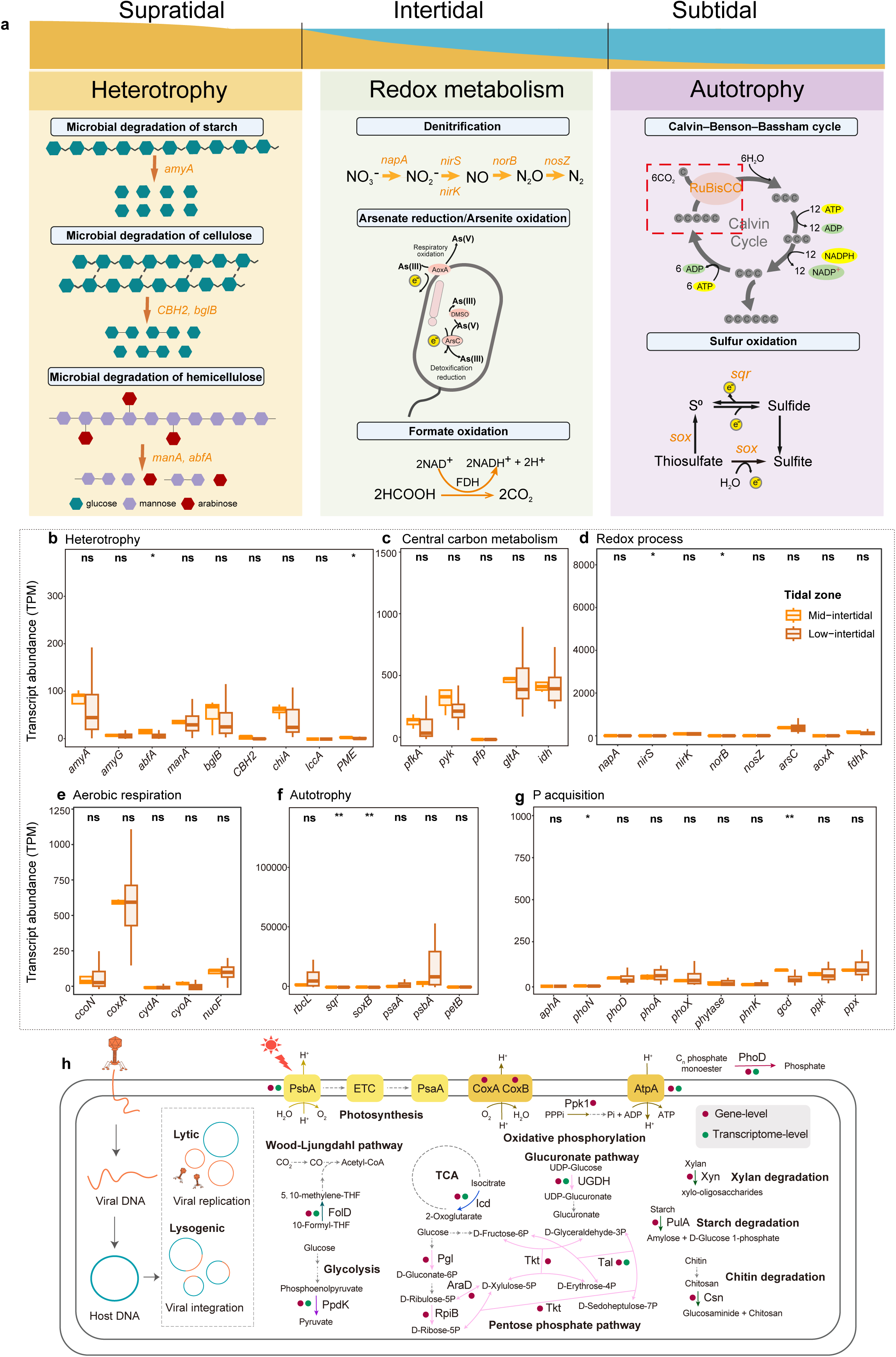
Metabolic differentiation of sandy beach microbial and viral communities across tidal zones. **(a)** Conceptual schematic summarizing metabolic zonation across sandy beach habitats, highlighting the predominance of heterotrophic processes in supratidal sediments and the increasing importance of redox metabolism and autotrophic pathways toward the subtidal zone. For summary visualization, habitats are grouped into supratidal, intertidal and subtidal zones. **(b–g)** Transcript abundances (TPM) of functional marker genes involved in heterotrophy **(b)**, central carbon metabolism **(c)**, redox processes **(d)**, aerobic respiration **(e)**, autotrophy **(f)** and phosphorus acquisition **(g)**. Transcripts were quantified from metatranscriptomes generated from water samples collected in the mid- and low-intertidal zones. Colors denote tidal position. Boxplots show medians (center lines), interquartile ranges (boxes) and 1.5 × interquartile-range whiskers. Statistical differences were evaluated using Kruskal–Wallis tests, with significance indicated as ns, \**P* < 0.05, \*\**P* < 0.01 and \*\*\**P* < 0.001. In panel b, PME denotes pectinesterase. **(h)** Schematic overview of viral auxiliary metabolic genes detected in sandy beach viromes, showing potential viral modulation of host pathways related to photosynthesis, oxidative phosphorylation, carbon metabolism, phosphorus metabolism and polysaccharide degradation. Dots indicate gene-level and transcriptome-level evidence, as shown in the panel.

Toward the intertidal zone, microbial communities displayed increasing reliance on redox-flexible metabolic pathways (**Fig. 5a**). Genes associated with alternative electron acceptors, including denitrification (*napA*, *nirS*, *nirK*, *norB* and *nosZ*; mean 16-124 GPM), arsenite oxidation (*aoxA*; mean 13 GPM), arsenate reduction genes (*arsC*; mean 722 GPM) and formate oxidation (*fdhA*; mean 376 GPM), were widely detected and tended to show relatively higher abundance in intertidal sediments (**Figs. S17c-d**).

These genes were mainly affiliated with *Pseudomonadota* and *Desulfobacterota* lineages, suggesting that these taxa contribute to respiration under fluctuating oxygen conditions (**Tables S5-S6**). In contrast, autotrophic and chemolithotrophic pathways became increasingly prominent toward the subtidal zone (**Fig. 5a**). RuBisCO genes (*rbcL*; mean 50 GPM), representing the Calvin–Benson–Bassham cycle, were significantly enriched in subtidal sediments (Kruskal–Wallis test, *P* < 0.001), while sulfur oxidation genes, including sulfide:quinone oxidoreductase (*sqr*; mean 155 GPM) and thiosulfate oxidation (*soxB*; mean 54 GPM), showed similar seaward enrichment (**Figs. S17f-g**). These pathways were predominantly associated with *Pseudomonadota* and *Campylobacterota*-related lineages, which include known chemolithoautotrophic microorganisms^49^. Phototrophic marker genes (*psaA* and *psbA*) were detectable but generally low in abundance, although relatively higher signals were observed in subtidal zones (**Fig. S17h**).

Phosphorus acquisition strategies were pervasive across all tidal zones (**Fig. S17e**). Alkaline phosphatase genes (*phoA*, *phoD* and *phoX*; mean 146 GPM), which mediate organic phosphorus mineralization, were substantially more abundant than acid phosphatases, while genes involved in mineral phosphate solubilization (*gcd*) and polyphosphate metabolism (*ppk* and *ppx*) were also widespread. These patterns indicate that microbial communities employ diverse strategies to access phosphorus under nutrient-limited conditions (**Table S6**), independent of tidal position^50^.

Metatranscriptomic analyses further showed that these pathways were actively expressed in water samples from the mid- and low-intertidal zones, with broadly similar expression patterns between the two zones (**Fig. 5b–g and Table S25**). Transcripts associated with carbon fixation (*rbcL*) and sulfur oxidation (*sqr*) were consistently abundant in intertidal waters, supporting the ecological importance of chemolithotrophic and autotrophic processes in these environments^7^. These results indicate that tidal zonation structures not only metabolic potential but also in situ metabolic activity.

Beyond cellular microorganisms, sandy beach viruses may further shape these metabolic processes through auxiliary metabolic genes (AMGs). Among 38,255 identified vOTUs, 4,641 AMGs (143 gene families encoded by 2,556 vOTUs) were detected that overlap with dominant host metabolic pathways, including photosynthesis (*psbA*), aerobic respiration (*coxA*, *coxB* and *atpA*), phosphorus metabolism (*phoD* and *ppk*) and carbon metabolism genes (e.g., *icd*, *ppdK*, *xyn*, *pulA, csn*) (**Fig. 5h and Table S26**). Several of these AMGs (n = 233, spanning 57 gene families; e.g., *psbA*, *atpA*, *phoD*, *icd*, *ppdK*) also exhibited detectable transcriptional activity, with nearly 40% of gene families being actively expressed, highlighting the widespread transcriptional engagement of viral AMGs. This indicates that viral infection can influence actively expressed pathways. The strong functional overlap between viral AMGs and host metabolic pathways suggests that viruses may modulate and potentially amplify key biogeochemical processes along the tidal gradient.

### Sandy beach microbiomes encode broad capacities for pollutant transformation and microbial chemical interactions

Hydrocarbon degradation genes were broadly represented, with 2,915 proteins spanning 24 functional categories covering pathways for short- and medium-chain alkanes, long-chain alkanes, monoaromatic hydrocarbons and polycyclic aromatic hydrocarbons (**Fig. 6a and Table S27**). Most sequences (2,841 genes; 97.5%) were associated with aerobic pathways, consistent with efficient oxygen renewal in permeable sediments^51–53^. Key enzymes included AlkB, CYP153, AlmA, LadA, MAHα/β and NdoB/NdoC, indicating capacity for oxidation of chemically diverse petroleum-derived substrates. Among these, AlkB (mean: 31 GPM) and CYP153 (mean: 37 GPM) were particularly abundant and showed significant variation across tidal zones (**Fig. 6b and Table S28**; *P* < 0.001). Moreover, these hydrocarbon-degradation genes were actively expressed in water samples from the mid- and low-intertidal zones, with CYP153 showing especially high expression (mean: 59 TPM; **Table S29**). Phylogenetic reconstruction resolved nine AlkB clades, including a previously unrecognized lineage lacking close reference homologues, and structural modelling showed strong similarity to experimentally validated AlkB enzymes (PDB: 8F6T; TM-score = 0.98) with conserved catalytic residues (**Figs. S18–S19**). Hydrocarbon degradation genes were detected in 1,606 MAGs, including bacterial lineages from *Gammaproteobacteria*, *Actinomycetota* and *Binatia*, as well as archaeal homologues affiliated with *Thermoplasmatota* and *Halobacteriota* (**Table S27**), extending the phylogenetic breadth of aerobic hydrocarbon oxidation^54–56^. In total, 38 MAGs encoded multiple hydrocarbon degradation modules, frequently combining *alkB*, *CYP153*, *MAHα/β* and *ndo* genes, with co-localized gene clusters suggesting coordinated oxidation of multiple hydrocarbon classes **(Fig. 6c and Fig. S20**).

**Figure 6.**
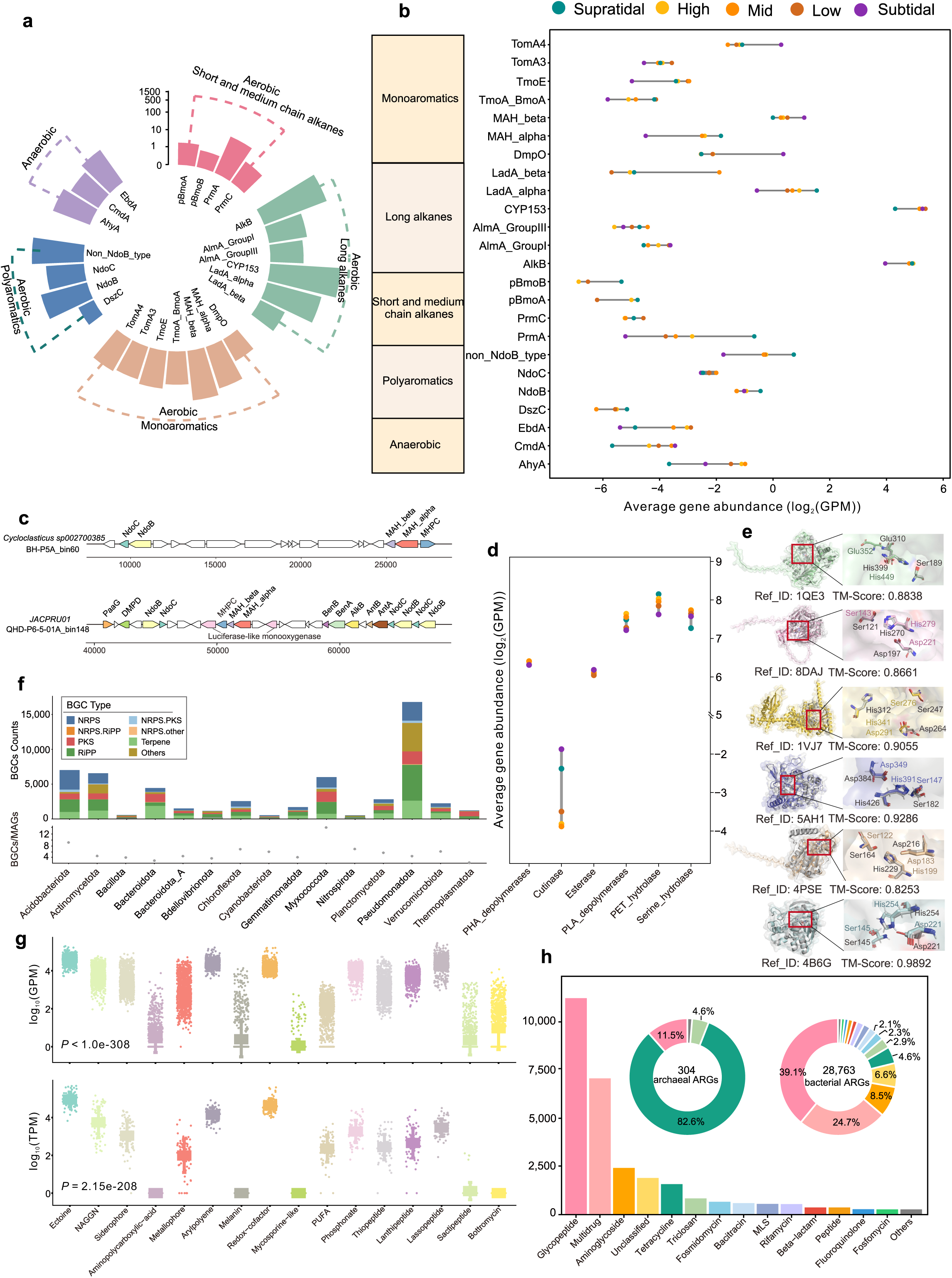
Sandy beach microbiomes encode pollutant transformation, secondary metabolism and antibiotic resistance functions. **(a)** Functional profile of hydrocarbon-degradation genes in the SMGC, grouped by substrate class and degradation mode. **(b)** Average abundance of hydrocarbon-degradation genes across tidal zones, expressed as log_2_-transformed genes per million (GPM). **(c)** Representative hydrocarbon-degradation gene clusters in two MAGs, illustrating co-localized pathways for oxidation of multiple hydrocarbon classes. **(d)** Average abundance of plastic-degrading enzyme classes across tidal zones, expressed as log_2_-transformed GPM. **(e)** Structural alignments of representative plastic-degrading enzymes with experimentally characterized reference proteins. Catalytic residues, reference identifiers and TM-scores are shown. **(f)** Distribution of biosynthetic gene cluster (BGC) classes across major bacterial and archaeal phyla in the SMGC. **(g)** Gene and transcript abundances of selected BGC-associated product classes and adaptive metabolites, expressed as log_10_-transformed GPM and TPM, respectively. **(h)** Composition of antibiotic resistance genes (ARGs) in the SMGC, including ARG counts by resistance class and the compositions of archaeal and bacterial ARG repertoires. Detailed annotations are provided in **Table S36**.

In parallel, the SMGC revealed an extensive repertoire of putative plastic-degrading enzymes, indicating a broader xenobiotic transformation network in sandy beach ecosystems (**Fig. S21 and Table S27**). We identified 11,929 polyethylene terephthalate (PET) hydrolases, 10,528 polylactic acid (PLA) depolymerases, 9,332 serine hydrolases, 5,583 polyhydroxyalkanoates (PHA) depolymerases, 4,765 esterases, and 45 cutinases, collectively targeting diverse polyester and aliphatic polymers^57^. These genes were widely distributed across 10,949 bacterial MAGs spanning 64 phyla, dominated by *Pseudomonadota*, *Actinomycetota* and *Bacteroidota*, with additional archaeal homologues (n = 327) primarily affiliated with *Thermoplasmatota*, indicating broad phylogenetic involvement in plastic transformation. Except for cutinases, all enzyme classes showed significant variation across tidal zones (**Fig. 6d and Table S30**; *P* < 0.001), with highest abundances generally observed in intertidal sediments (e.g., PET hydrolases mean 252 GPM), consistent with enhanced accumulation of microplastics and organic substrates at the land-sea interface^58^. Furthermore, intertidal water metatranscriptomes confirmed *in situ* expression of the plastic-degrading genes (e.g., PET hydrolases, mean 84 TPM; **Table S31**). Structural prediction of representative enzymes yielded TM-scores of 0.82–0.98 (**Fig. 6e**), revealing conserved α/β-hydrolase folds and catalytic residues characteristic of functional polyester hydrolases^59, 60^. Notably, plastic- and hydrocarbon-degrading genes frequently co-occurred within the same genomes (**Table S27**), indicating multi-substrate metabolic versatility and suggesting adaptation to chemically heterogeneous pollutant mixtures shaped by tidal transport and retention^61, 62^.

Beyond xenobiotic degradation, sandy beach microbiomes encoded extensive biosynthetic capacities, shaping microbial chemical ecology. Across the SMGC, we identified 58,956 biosynthetic gene clusters (BGCs), including 57,546 bacterial and 1,410 archaeal clusters, spanning 29 biosynthetic classes, including both canonical and hybrid pathways (**Figs. S22–S23 and Table S32**). The dominant BGC categories included RiPPs (14,842 clusters), NRPS (12,288), terpenes (11,845), and PKSs (8,873), together accounting for more than 80% of bacterial biosynthetic diversity. Near-complete genomes (completeness ≥90%, contamination ≤5%) and BGC-rich (≥15 BGCs) genomes (n = 111) also contained extracellular polysaccharide (exoPS) biosynthesis clusters, of which 81% corresponded to uncharacterised Wzy-dependent or ABC transporter-dependent pathways (**Table S33**), particularly in *Acidobacteriota* and *Myxococcota*, potentially supporting adhesion, aggregation and biofilm formation in dynamic sediments^63^. BGC distribution largely followed community composition (**Fig. 6f**), with major contributions from *Pseudomonadota*, *Actinomycetota*, *Acidobacteriota* and *Myxococcota*, the latter showing the highest biosynthetic density (up to 14 BGCs per genome), consistent with strong metabolic specialization.

Functionally, many BGCs were associated with adaptation to environmental stress characteristic of sandy beaches (**Fig. 6g and Table S34**). Osmoprotectant pathways such as ectoine^64^ (mean 40,059 GPM) and N-acetylglutaminylglutamine amide^65^ (NAGGN; mean 10,412 GPM) were highly abundant, consistent with fluctuating salinity and recurrent wetting-drying cycles, while pigment-associated arylpolyene clusters^66^ (25,874 GPM) and redox cofactor pathways (mean 15,569 GPM) support tolerance to oxidative and UV stress. Metal acquisition pathways, including siderophores (mean 5,995 GPM) and metallophores (mean 3,341 GPM), indicate adaptation to nutrient limitation in permeable sands, and polyunsaturated fatty acid pathways (mean 211 GPM) suggest membrane-level responses to environmental variability. Antimicrobial-associated BGCs were also prevalent, including lanthipeptide, lassopeptide and thiopeptide pathways^67^, with lassopeptides (mean 31,173 GPM) representing the most abundant antimicrobial class, together with phosphonate pathways (mean 13,784 GPM). Metatranscriptomic analyses further showed that stress-adaptation BGCs exhibited higher expression than antimicrobial clusters (**Fig. 6g and Table S35**; *P* < 0.001), with particularly high transcript abundance for ectoine (mean 129,151 TPM), NAGGN (mean 20,395 TPM) and arylpolyene (mean 19,351 TPM) pathways, indicating strong selection imposed by salinity fluctuation, desiccation-rewetting dynamics and oxidative stress.

Microbial chemical defense capacity was further reflected in the extensive antibiotic resistome (**Fig. 6h**). The SMGC contained 29,067 antibiotic resistance genes (ARGs) identified in 10,505 MAGs spanning 62 phyla, indicating widespread adaptation to bioactive compounds derived from both microbial competition and anthropogenic inputs (**Table S36**). Archaeal genomes encoded 304 ARGs, whereas bacterial genomes carried 28,763 ARGs across 28 resistance classes, dominated by glycopeptide (39.1%), multidrug (24.7%) and aminoglycoside (8.5%) resistance genes. ARGs frequently co-occurred with BGCs within the same genomes (**Table S37**), particularly in *Actinomycetota* (1,163 MAGs) and *Pseudomonadota* (3,137 MAGs), indicating tight evolutionary coupling between secondary metabolite production and resistance capacity^68^. Environmental measurements further detected 12 antibiotics in 276 sand samples and 14 antibiotics in 54 water samples (**Table S7**), including azithromycin (up to 101 pg g^-1^) and sulfaquinoxaline (up to 60 pg g^-1^) in sand, consistent with persistent exposure to antimicrobial compounds.

## Discussion

Sandy beaches are physically dynamic environments in which hydrodynamic forcing strongly shapes sediment structure, oxygen availability and solute transport. Across a continental-scale dataset spanning diverse climatic and morphodynamic settings, tidal elevation consistently structured microbial diversity, metabolic organization and virus-host interactions. This repeatable zonation pattern indicates that cross-shore position provides a primary constraint on microbiome assembly in permeable sediments, outweighing the effects of geographic variation and differences in beach usage detected in this dataset^7, 15, 17^. Because inundation frequency controls oxygen exposure, porewater residence time and resource renewal, tidal forcing generates predictable environmental gradients that organize microbial community composition along the land-sea interface. The metabolic profiles observed across tidal zones are consistent with these physical constraints. Upper beach sediments were dominated by aerobic heterotrophic pathways associated with rapid turnover of labile organic matter, whereas sediments toward the subtidal zone showed increasing representation of chemolithotrophic and autotrophic processes linked to fluctuating redox conditions. Permeable sands differ from diffusion-dominated sediments in that advective exchange rapidly redistributes electron donors and acceptors, creating transient oxic-anoxic interfaces at millimetre scales^17, 69^. Such conditions favor metabolically flexible microorganisms capable of switching between respiratory strategies, explaining the coexistence of aerobic respiration, denitrification and sulfur oxidation pathways across short spatial distances.

At the genomic scale, sandy beach sediments harbor exceptionally high diversity and novelty, consistent with a distinct evolutionary reservoir rather than a simple admixture of marine and terrestrial microorganisms. The limited overlap between the SMGC and existing environmental genome collections indicates that permeable sediments support specialized lineages adapted to dynamic microscale gradients, repeated disturbance and pulsed resource supply. Elevated novelty in mid-intertidal sediments further supports the idea that fluctuating ecotones promote diversification by generating heterogeneous niches and episodic selection pressures^11, 14, 70^. Viral communities exhibit similarly high levels of novelty and habitat specificity, with limited overlap with global virome datasets. The strong correspondence between viral assemblages and host distributions, together with extensive virus-host linkages and auxiliary metabolic genes, supports the view that viruses are a major component of sandy beach ecosystems, potentially influencing microbial turnover, metabolic evolution and biogeochemical cycling^71, 72^.

In addition to core carbon and nutrient cycling, sandy beach microbiomes encode broad potential for hydrocarbon and plastic transformation. The predominance of aerobic hydrocarbon degradation genes is consistent with efficient oxygen renewal driven by tidal pumping, while the widespread occurrence of candidate polyester hydrolases is consistent with adaptation to recurrent exposure to synthetic polymers accumulating at the land-sea interface^73–75^. The frequent co-occurrence of hydrocarbon and plastic degradation genes within the same genomes is consistent with metabolic versatility under chemically mixed conditions. These traits are consistent with the role of permeable sediments as zones of intense solute exchange, where both natural organic matter and anthropogenic compounds are repeatedly transported and retained. Biosynthetic gene clusters and antibiotic resistance genes were also widely distributed, reflecting the importance of both abiotic stress tolerance and microbial interactions in sandy sediments. High abundances of osmoprotectant and oxidative stress-related pathways indicate adaptation to fluctuating salinity, desiccation-rewetting cycles and strong irradiance. The co-occurrence of biosynthetic and resistance genes within the same genomes suggests a potential linkage between secondary metabolite production and resistance capacity, consistent with competitive interactions in spatially structured microbial communities. The detection of diverse antibiotic residues in sand and water further indicates that microbial communities are recurrently exposed to bioactive compounds derived from both natural processes and human activities^18, 38, 76^.

Together, these results indicate that sandy beaches represent physically structured microbial ecosystems characterized by high genomic novelty, strong metabolic flexibility and broad functional potential for processing chemically diverse substrates. By integrating cross-shore sampling with genome-resolved multi-omics analyses, this study provides a reference framework for investigating microbiome dynamics in permeable coastal sediments and for assessing how physical forcing and environmental variability jointly shape microbial community function in permeable coastal sediments.

## Materials and Methods

### Study sites and sampling design

We conducted a coordinated coast-wide survey of sandy beaches along the Chinese coastline during August-September 2024, corresponding to the summer tourism season. The survey spanned a latitudinal gradient from 18° N to 40° N and covered tropical, subtropical and temperate climatic zones across the South China Sea, East China Sea, Yellow Sea, and Bohai Sea. In total, 18 sandy beaches were investigated, including 15 urbanized recreational beaches and three natural reference beaches distributed across temperate, subtropical and tropical regions, respectively (**Fig. 1 and Table S1**). Sampling sites were selected to ensure broad geographic coverage of China’s coastline, representation of beaches experiencing high human activity, inclusion of natural reference beaches for baseline comparison and coverage of multiple beach morphodynamic types including reflective, intermediate and dissipative beaches. At each beach, depending on beach width / accessibility / tidal exposure, 7-10 cross-shore transects were established extending from the supratidal zone through the high-, mid-and low-intertidal zones, resulting in 174 standardized transects. Because the lower intertidal zone was accessible only during brief periods around extreme low tide, sampling in this zone was temporally constrained and yielded fewer samples.

### Field sampling and sample processing

Sediment sampling was conducted during low tide along each transect. Surface sediments (0–5 cm depth) were collected using metal-free polypropylene scrapers after removing visible debris such as shell fragments and plant material. Samples for omics analyses were transferred into sterile polypropylene tubes or pre-combusted aluminum containers, flash-frozen in liquid nitrogen *in-situ* and stored at −80 °C until nucleic acid extraction. Sediment samples for physicochemical analyses were transported under cooled and dark conditions, air-dried and sieved in the laboratory prior to analysis.

Water samples were collected adjacent to each transect using pre-cleaned polycarbonate or glass sampling bottles. For microbial metagenomic and metatranscriptomic analyses, water samples were sequentially prefiltered through 200 μm and 20 μm meshes to remove large debris and plankton and subsequently filtered through 0.22 μm polyethersulfone membranes to capture microbial cells. For viral metagenomes and viral metatranscriptomes, water samples were prefiltered using the same meshes and then filtered through 0.1 μm polyethersulfone membranes under low vacuum pressure. Filters were immediately frozen in liquid nitrogen and stored at −80 °C until nucleic acid extraction. For chemical analyses, water samples were filtered through 0.45 μm membranes to obtain the dissolved fraction. Samples for metal analysis were acidified to pH <2 using ultrapure HNO_3_ (65–69%, trace-metal grade) and stored at 4 °C in the dark. Nutrient samples were refrigerated and analyzed within 24 h, whereas antibiotic samples were frozen immediately after filtration.

### Morphodynamic characterization

Cross-shore transects were surveyed using a STONEX S9II PRO RTK-GPS system with a horizontal accuracy of ±8 mm and vertical accuracy of ±15 mm. Beach morphology was quantified using beach slope, dry-beach width and intertidal width. Based on these characteristics and surf-zone features, beaches were classified as reflective, intermediate or dissipative systems. Hydrodynamic conditions including tides and waves were characterized using field observations, nearby instrumental records and numerical simulations. Wave parameters including significant wave height, wave period and dominant wave direction were simulated using the TOMAWAC wave model^77^. Bathymetry was derived from the GEBCO global dataset combined with CMAP coastal bathymetric data. Sediment grain-size distributions were measured using dry sieving with a POWTEQ SS2000 vibrating sieve shaker after oven drying samples at 60 °C.

### Physicochemical analyses

In situ water physicochemical parameters including temperature and pH were measured using a YSI ProDSS multiparameter probe calibrated prior to deployment. Nutrient samples were collected in 500-mL polyethylene (PE) bottles and frozen before measurement. Dissolved inorganic nitrogen (nitrate, nitrite, and ammonium) and phosphate concentrations were analyzed using a 723C spectrophotometer (Shanghaijingke®, China) following standardized colorimetric methods. Ammonium was measured using the Nessler reagent colorimetric method (HJ 535-2009). Total nitrogen was determined after alkaline persulfate digestion by UV spectrophotometry (HJ 636-2012). Total phosphorus was measured after digestion using the ammonium molybdate spectrophotometric method (GB/T 11893-1989). Dissolved silicate was analyzed using a standard molybdenum blue spectrophotometric method. Metal concentrations were determined using inductively coupled plasma mass spectrometry (ICP-MS; Agilent 7900). Water samples were filtered through 0.45 μm membranes and acidified prior to analysis. Sediment samples were digested using mixed-acid digestion (HNO_3_-HCl-HF) in a microwave digestion system and subsequently analyzed by ICP-MS. Antibiotics in water samples were extracted using solid-phase extraction with Oasis HLB cartridges and quantified using liquid chromatography-tandem mass spectrometry (UPLC-MS/MS; Thermo TSQ) operating in multiple reaction monitoring (MRM) mode. Sediment antibiotics were extracted from freeze-dried sediments using methanol-dichloromethane mixtures followed by SPE purification prior to UPLC-MS/MS analysis.

### DNA extraction and metagenomic sequencing

Metagenomic DNA was extracted from 978 sand and water samples. For water metagenomes, the cellular fraction was collected on 0.22 μm filters prior to DNA extraction. For an additional 63 water samples used for virome analysis, seawater was first filtered through 0.22 μm membranes to remove cells, amended with 0.1% ferric chloride for 30 min to induce viral flocculation, and then filtered to recover virus-enriched material for DNA extraction. To optimize DNA recovery from samples with different moisture conditions, moist samples (>20% moisture content) were extracted using the E.Z.N.A. Soil DNA Kit (Omega Bio-Tek, Norcross, GA, USA), whereas drier samples were extracted using the DNeasy PowerSoil Pro Kit (QIAGEN, Netherlands). Samples yielding <500 ng DNA were re-extracted and pooled to obtain at least 1 μg total DNA. DNA concentration and purity were evaluated spectrophotometrically. Samples with OD_260/280_ ratios of 1.8–2.2 and OD_260/230_ values ≥2.0 were used for library preparation. Sequencing libraries were prepared using a standard genomic DNA library preparation procedure workflow and sequenced by Biozeron Biological Technology Co., Ltd. (Shanghai, China) on the MGI-T7 platform in paired-end mode (2 × 150 bp).

### Community profiling, metagenomic assembly and genome reconstruction

Raw reads were quality-filtered and adaptor-trimmed using Trimmomatic (v0.39)^78^. Clean reads were then profiled using sylph (v0.8.0)^79^ with default parameters against the GTDB r220-c200 and IMG/VR c200 reference databases to characterize the composition of prokaryotic and viral communities, respectively; prokaryotic profiling was performed on 978 samples, whereas viral profiling included these samples together with an additional 63 virome samples. Clean reads were also assembled using MEGAHIT (v1.1)^80^ with the parameter “--min-contig-len 500”. Contig coverage was calculated using jgi_summarize_bam_contig_depths from MetaBAT2 (v2.12.1)^81^ based on sorted BAM files generated with BWA-MEM (v0.7.17)^82^ and SAMtools (v1.21)^83^. Contigs were binned using MetaBAT2 (v2.12.1) and SemiBin2 (v 2.1.0)^84^. Bins generated by the two methods were refined using the bin_refinement module in MetaWRAP (v1.3.2; parameters: -c 50 -x 10)^85^. Species-level dereplication was performed using dRep (v3.5.0; parameters: -comp 50 -con 10 -sa 0.95 --S_algorithm skani)^86^. Genome quality was assessed using CheckM2 (v1.1.0)^87^. MAGs were retained if they met the criterion of medium or higher quality of the MIMAG standard (completeness ≥50%, contamination ≤10%, and quality score QS = completeness − 5 × contamination ≥50). In total, 13,337 MAGs were retained for downstream analyses and designated as the SMGC.

Taxonomic annotation of MAGs was performed using GTDB-Tk (v2.4.1)^88^ against GTDB release 226. Phylogenetic diversity within each phylum was assessed using GenomeTreeTk (v0.1.6; https://github.com/dparks1134/GenomeTreeTk). Relative abundances of MAGs were calculated using CoverM (v0.7.0)^89^ with reads-per-kilobase-million (RPKM) normalization. Comparative analyses were performed between the SMGC and published genome catalogues from other ecosystems, including the Global Ocean Microbiome Catalogue (GOMC)^29^, the Soil Microbiome Catalogue^28^, the Genome-Resolved Open Watersheds Database (GROWdb)^27^, the Mariana Trench Environment and Ecology Research dataset (MEER)^30^ and the Global Cold Seep Microbiome Catalogue^46^.

### Viral genome identification and characterization

Potential viral sequences were identified from assembled contigs across 978 metagenomes and 63 viromes using geNomad (v1.8.1; end-to-end) ^90^ and VirSorter2 (v2.2.4; default parameters)^91^. Viral completeness and contamination were evaluated using CheckV (v1.0.3; database v1.5)^92^, and contigs with estimated completeness ≥50% were retained. Viral contigs were clustered into species-level vOTUs using a 95% average nucleotide identity threshold and 85% minimum aligned fraction, following CheckV recommendations. This yielded 38,255 vOTUs, collectively defined as the SVC. Taxonomic assignments were generated using geNomad (v1.8.1)^90^ against ICTV release MSL39. Open reading frames were predicted using Prodigal (v2.6.3; -p meta) and functionally annotated using DRAM-v (v1.3.5)^93^. Major capsid proteins were identified by searching predicted viral proteins against the PDB100 database^94^ using Foldseek^95^ in easy-search mode with an E-value cutoff of 1 × 10^-5^. Auxiliary metabolic genes were inferred from DRAM-v annotations, and functional filters were applied to obtain high-confidence AMGs. Viral lifestyle was predicted using phabox (v2.1.13) with the phatyp task^39^. Viral novelty and environmental distribution were assessed by comparing SVC vOTUs with published virome catalogues from mudflat intertidal sediments^23^ (n = 20,102), soils^36^ (n = 80,750), groundwater^35^ (n = 280,420), the Global Ocean Viromes 2.0^34^ (GOV2, n = 488,133), and oil reservoirs^37^ (n = 3,886)^34^ using BLASTn (v2.16.0; -perc_identity 90 -qcov_hsp_perc 90)^96^.

### Virus-host association prediction

Virus-host associations were predicted using iPHoP (v1.4.1)^42^ with the iPHoP reference database and the 13,337 MAGs recovered in this study. The pipeline integrated sequence-based and composition-based approaches, including BLASTn searches against host and CRISPR spacer databases and host prediction by RaFAH (v0.3)^97^, WIsH (v1.0)^98^, VirHostMatcher-Net^99^, and PHP^100^. Virus-host associations supported by multiple independent lines of evidence were considered high-confidence, with CRISPR spacer matches treated particularly strong evidence. In cases where multiple candidate hosts were detected, the host with the highest confidence score was selected as the final assignment.

### Gene prediction and functional annotation

Protein-coding sequences were predicted from assembled contigs >1,000 bp using Prodigal (v2.6.3; -meta)^101^. The resulting sequences were clustered hierarchically at 90% amino acid identity and 90% coverage using MMseqs2 v13.45111 with the easy-linclust module (-c 0.9 -min-seq-id 0.9 --cov-mode 1)^102^. This generated a non-redundant gene catalogue comprising 407,984,259 representative clusters. Representative amino acid sequences were functionally annotated using eggNOG-mapper (v2.1.12)^103, 104^. Functional assignments included eggNOG 5.0, Pfam 33.1, KEGG, EC, GO and CAZy.

Protein-coding sequences of genomes in SMGC were predicted with Prodigal (v2.6.3) in single mode, and then annotated with eggNOG-mapper (v2.1.12) following the same procedure as described above. Annotated KO numbers were used for inferring the metabolic pathway encoded in each MAG. Hydrocarbon degradation genes were identified using hmmsearch against 37 HMM profiles with noise cutoff from the CANT-HYD database^105^. Plastic-degrading proteins were identified using hmmsearch against the RemeDB database with default parameters^106^. Secondary metabolite BGCs were identified using antiSMASH (v7.1)^107^ with the parameters “-fullhmmer -cb-general - cb-subclusters -cb-knownclusters -genefinding-tool prodigal-m -clusterhmmer -asf - smcog-trees -pfam2go”. Among SMGC genomes, 111 BGC-rich MAGs (≥15 BGCs; completeness ≥90%, contamination ≤5%) were selected for exopolysaccharide BGC prediction using epsSMASH (v1.0)^108^ with --genefinding-tool prodigal. ARGs were inferred using RGI v6.0.3 in strict mode with CARD (v3.2.4)^109^ and independently annotated using DeepARG v2 in long-sequence mode (--min-prob 0.8 --id 50 -e 1e-10)^110^. DeepARG results were normalized to ARO terms using ArgNorm (v1.0.0)^111^. A sequence was retained as an ARG if it was identified by either approach. Putative dehalogenases annotated by eggNOG-mapper were further validated by BLASTp against UniProt. Only sequences with ≥60% identity, ≥90% query coverage and E-values <1 × 10^-5^ were retained^112^. Polyphenol-related genes were annotated using CAMPER v1.0.13 with default parameters^113^.

To quantify the abundance of functional genes, predicted proteins from SMGC genomes were clustered at 95% amino acid identity using CD-HIT (v4.8.1; -c 0.95 -aS 0.9 -g 1 - d 0)^114^. Metagenomic clean reads were then mapped to the non-redundant MAG gene catalogue with Salmon (v1.10.3; -validateMappings -meta)^115^ in mapping-based mode, and gene abundance was expressed as genes per million (GPM). For each BGC, the metagenomic abundance was estimated as the abundance of the cluster’s core gene sequence, quantified with the same Salmon workflow and reported in GPM.

### Phylogenetic analyses

Phylogenomic trees of bacterial and archaeal genomes were constructed using concatenated alignments of 120 bacterial and 53 archaeal conserved marker genes identified using GTDB-Tk (v2.4.1)^116^. Aligned marker genes were concatenated to generate genome-scale alignments and phylogenetic trees were inferred using FastTree^117^ (v2.1.11) under default parameters. Protein phylogenetic trees were constructed for selected functional genes (e.g., AlkB). Reference sequences of AlkB were retrieved from KEGG and UniProt databases. Protein sequences were aligned and phylogenetic inference was performed using IQ-TREE (v2.3.3)^118^ with model selection (−m MFP) and branch support estimated using 1000 ultrafast bootstrap replicates and SH-aLRT tests. All phylogenetic trees were visualized using iTOL (v7)^119^.

### Protein structure prediction and structural comparison

Representative protein sequences of selected functional genes (e.g., AlkB and plastic-degrading enzymes) were subjected to structural prediction using AlphaFold3 (https://alphafoldserver.com/)^120^. Predicted protein structures were evaluated using the predicted template modeling (pTM) score, and representative models were selected for structural comparison. Pairwise structural alignments between predicted structures and experimentally resolved reference proteins were performed using TM-align (v20170708; default parameters)^121^, and structural similarity was evaluated using TM-scores. Protein structures and alignments were visualized using PyMOL (v3.0.0)^122^.

### RNA extraction and metatranscriptomic sequencing

Total RNA was extracted using the RNeasy PowerSoil Total RNA Kit (QIAGEN) from 72 water samples, including 46 microbial metatranscriptome samples collected on 0.22 μm filters and 26 additional virus-enriched samples obtained by pre-filtration through 0.22 μm membranes, ferric chloride flocculation (0.1%, 30 min), and subsequent recovery of virus-enriched material prior to RNA extraction. Residual genomic DNA was removed by DNase treatment. Ribosomal RNA was depleted using a probe-based magnetic bead depletion workflow. The enriched RNA was reverse-transcribed, and cDNA libraries were prepared using the NEBNext Ultra II RNA Library Prep Kit following an Illumina-compatible protocol. Libraries were sequenced by Biozeron Biological Technology Co., Ltd. on the MGI-T7 platform in paired-end mode (2 × 150 bp).

### Metatranscriptomic quantification of microbial and viral activity

Raw metatranscriptomic reads were quality-filtered using the Read_QC module of MetaWRAP (v1.3.2; -skip-bmtagger)^85^. Ribosomal RNA sequences were removed using SortMeRNA (v4.3.7) with default settings^123^. Transcript abundances of prokaryotic functional genes and BGCs were determined by mapping clean reads to the non-redundant MAG gene catalogue and core biosynthetic genes using Salmon (v1.10.3; -validateMappings -meta)^115^. Gene transcript abundance was expressed as transcripts per million (TPM). For viral transcriptional activity, metatranscriptomic reads were mapped to vOTU sequences using Bowtie2 (v2.5.4) with default parameters^124^. SAM files were converted to BAM, sorted and indexed with SAMtools (v1.20). Alignments were filtered with CoverM (v0.7.0)^89^, retaining only reads aligned over ≥90% of their length with ≥95% nucleotide identity, as in a previous study^125^. Coverage profiles of viral contigs were generated using BEDTools genomecov (v2.31.1)^126^. To minimize spurious detection, only viral contigs with mapped reads covering at least 70% of the contig length were retained. Average gene-level coverage was then calculated using BEDTools coverage based on the filtered BAM files and Prodigal GFF annotations. A viral population was considered transcriptionally detected if at least one gene per 10 kb of viral genome had positive average coverage following a previous study^125^. Final vOTU coverage was calculated as the mean coverage of all genes within each viral population. Viral populations not meeting these detection criteria were assigned zero coverage. Gene-level coverage values were linked to functional annotations to assess AMG transcription.

### Statistical analysis

Statistical analyses were performed in R (v4.2.3). Alpha diversity was calculated using the Simpson index and compared among groups using Kruskal–Wallis tests, followed by pairwise two-sided Wilcoxon rank-sum tests where appropriate. Gene abundance and transcript abundance were compared among groups using the same approach. Community dissimilarities were calculated using Bray–Curtis distances and visualized by non-metric multidimensional scaling (NMDS) implemented in the vegan package. Differences in community composition were assessed using analysis of similarities (ANOSIM) with 999 permutations. Associations between environmental variables and community composition were assessed using Mantel tests. Unless otherwise stated, statistical significance was defined as *P* < 0.05.

## Data availability

All metagenomic, metatranscriptomic and viromic sequencing data generated in this study have been deposited in the National Genomics Data Center (NGDC) under accession number PRJCA0589xx. The genomic sequences of the SMGC and SVC are publicly available at https://doi.org/10.5281/zenodo.189340xx.

## Code availability

The present study did not generate codes, and the mentioned tools used for the data analysis were applied with default parameters unless specified otherwise.

## Acknowledgements

This work was supported by the National Key Research and Development Program of China (2022YFC3106100), the National Natural Science Foundation of China (U22A2058), and the Scientific Research Foundation of the Third Institute of Oceanography, Ministry of Natural Resources (2025035). We thank Jixiang Zheng for coordination of field campaigns and Haitao Xu for field safety oversight. We are grateful to colleagues who assisted with field sampling and logistical support, including Chengtao Wang, Zheyu Xiao, Jiahao Liu, Zhanyang Qi, Enquan Zhang, Chengpeng Li, Zehao Wang, Tengkai Chen, Weiming Wu, Zhuoyun Chen, Rupeng Du, Yongxin Shen, Zhipeng Zhu and Xiyu Pu.

## Author contributions

F.C. conceived the study, supervised the project, secured funding and contributed to manuscript writing. X.D. contributed to study design, performed analyses and led manuscript writing. Y.H. and C.Z. conducted the multi-omics analyses and contributed to figure preparation and manuscript writing. H.Q. led the geological characterization and morphodynamic analyses. L.W. and Z.P. conducted the physicochemical measurements. Y.C., Zr.L., Zj.L. and X.G. assisted with multi-omics analyses and figure preparation. S.Z., J.L., S.L., C.R. and L.Z. contributed to data interpretation and manuscript revision. All authors discussed the results and approved the final manuscript.

