## Supplementary Figures for "Continental-scale multi-omics reveals a distinct microbial-viral biome in sandy beach ecosystems structured by tidal zonation"

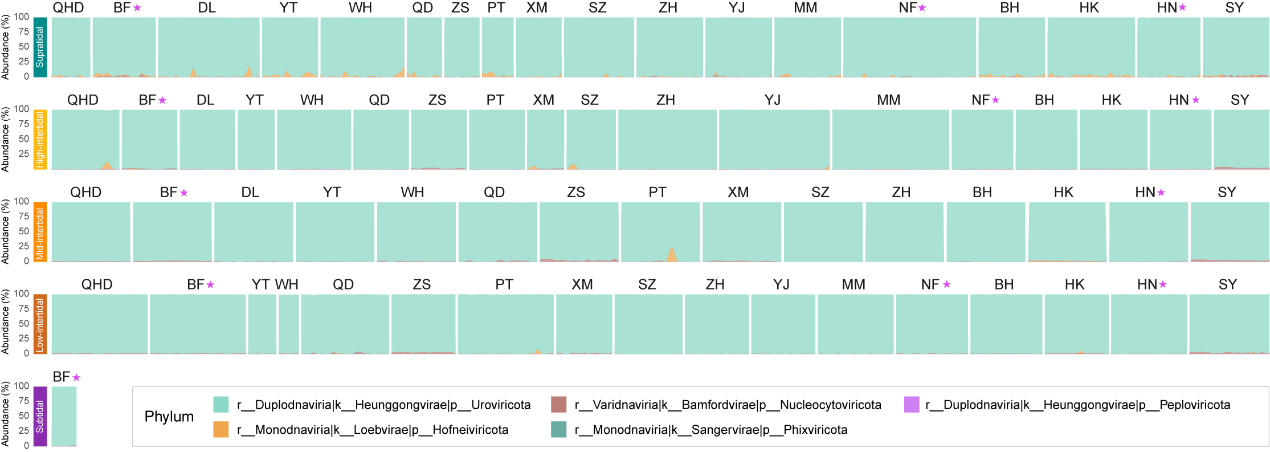

**Figure S1. Phylum-level taxonomic composition of viral communities across sandy beach sites.** Stacked bar plots show the relative abundance of viral taxa at the phylum level for each beach. Sites are ordered latitudinally from north to south to illustrate large-scale geographic trends in community composition. Colors denote different viral phyla. Asterisks indicate natural (low-disturbance) beaches. Detailed taxonomic information is provided in **Table S9**.

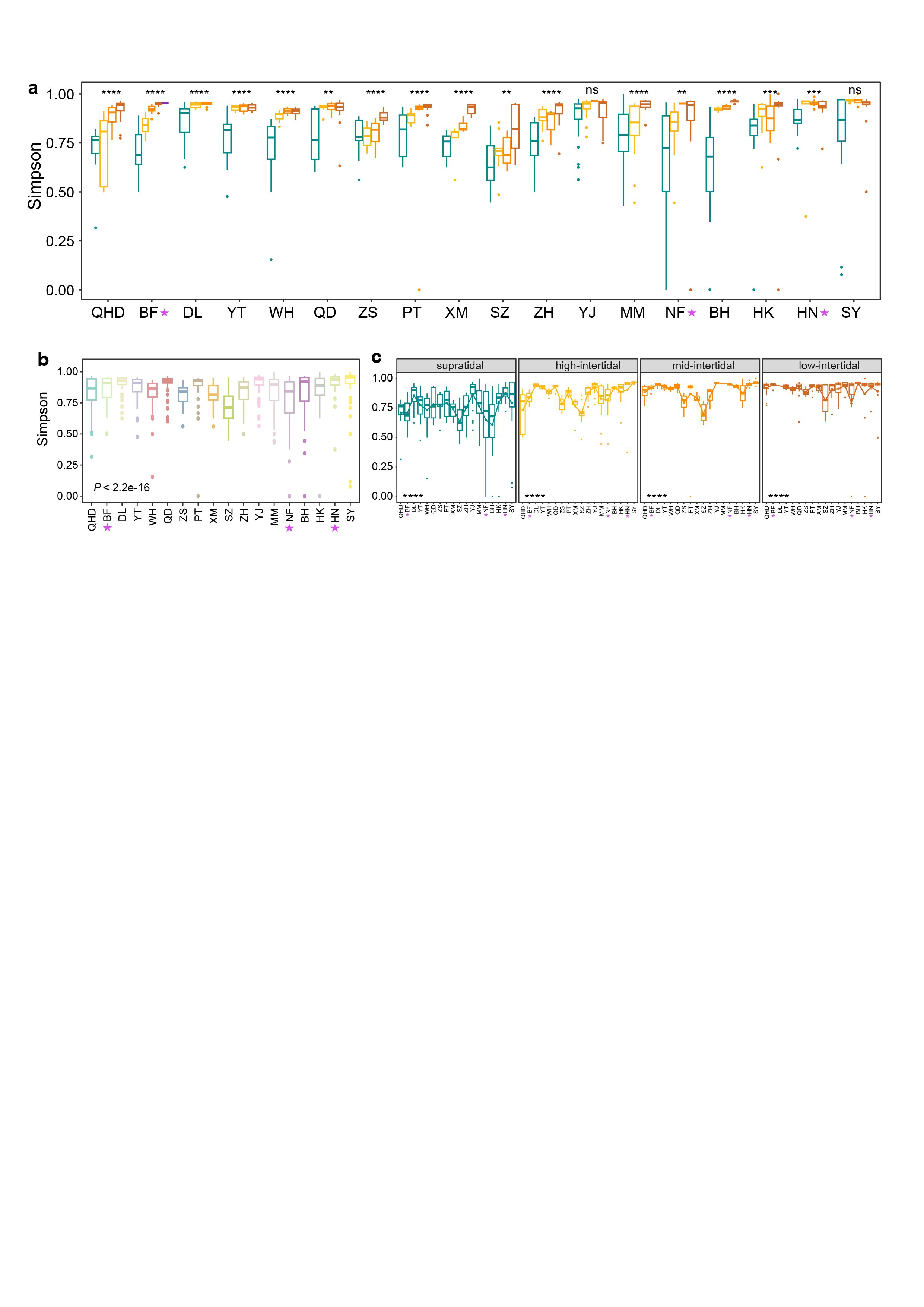

**Figure S2. Prokaryotic α-diversity across sandy beaches and tidal zones. (a)** Simpson diversity index compared among tidal zones within each beach. **(b)** Simpson diversity index grouped by beach and ordered latitudinally from north to south. **(c)** Simpson diversity index across beaches within each tidal zone (supratidal, high-, mid-, and low-intertidal). Boxplots show the median (center line), interquartile range (box), 1.5× interquartile range (whiskers), and outliers (points). Colors denote tidal zones. Statistical differences were evaluated using Kruskal–Wallis tests, with significance indicated as ns, *, **, and ***. Detailed statistics are provided in **Table S11**.

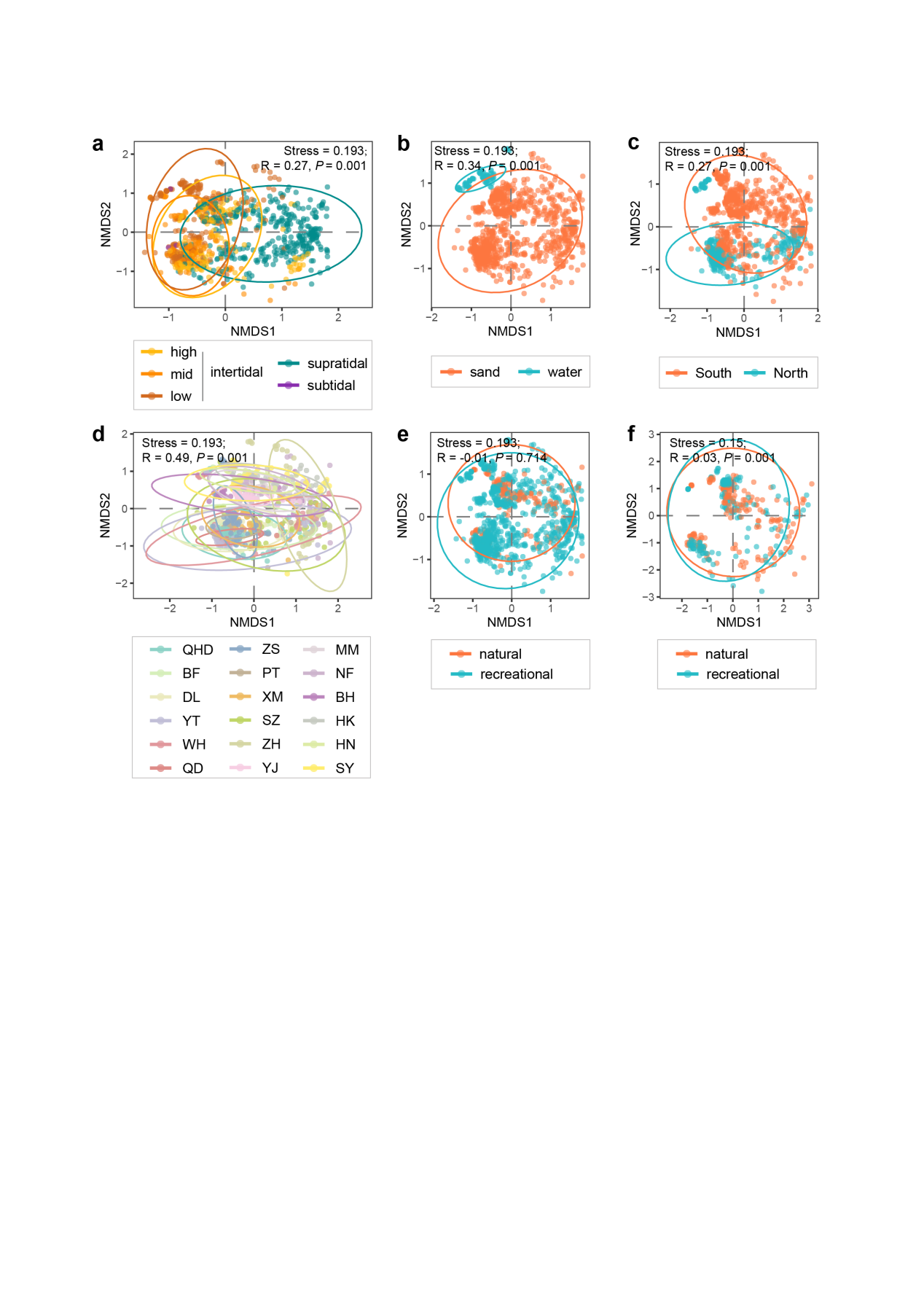

**Figure S3. Prokaryotic β-diversity across sandy beaches under multiple grouping schemes.** Non-metric multidimensional scaling (NMDS) ordinations based on Bray–Curtis dissimilarities showing community composition grouped by **(a)** tidal zones, **(b)** matrix type (sand versus water), **(c)** geographic regions (south versus north), **(d)** individual beaches (18 sites), and **(e–f)** beach usage (natural versus recreational). Panel **(e)** includes all samples, whereas panel **(f)** includes only paired recreational and adjacent natural beaches. Community dissimilarities were evaluated using ANOSIM with 999 permutations. Stress values, R statistics, and *P* values are shown in each panel.

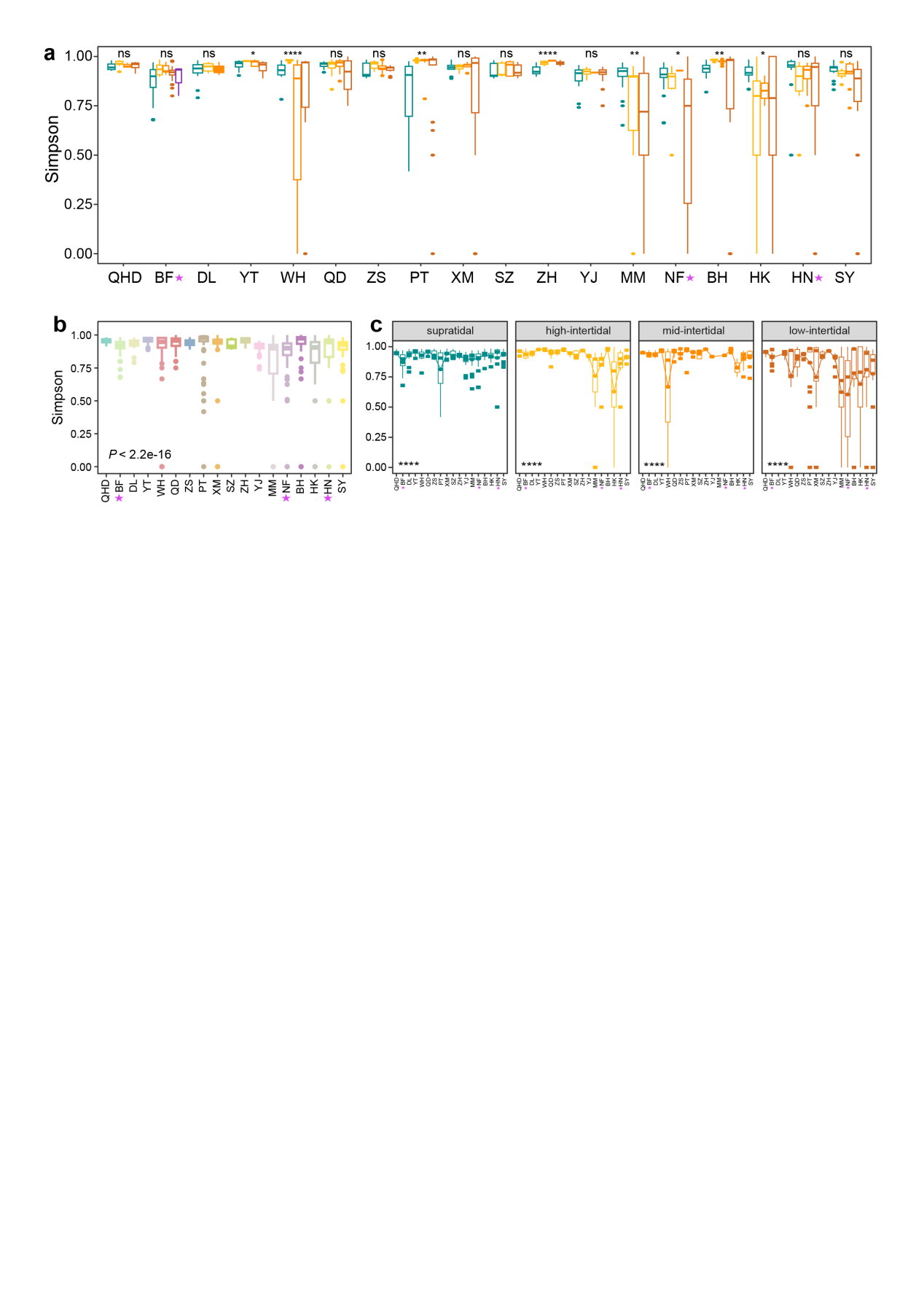

**Figure S4. Viral α-diversity across sandy beaches and tidal zones. (a)** Simpson diversity index compared among tidal zones within each beach. **(b)** Simpson diversity index grouped by beach and ordered latitudinally from north to south. **(c)** Simpson diversity index across beaches within each tidal zone (supratidal, high-, mid-, and low-intertidal). Boxplots show the median (center line), interquartile range (box), 1.5× interquartile range (whiskers), and outliers (points). Colors denote tidal zones. Statistical differences were evaluated using Kruskal–Wallis tests, with significance indicated as ns, *, **, and ***. Detailed statistics are provided in **Table S13**.

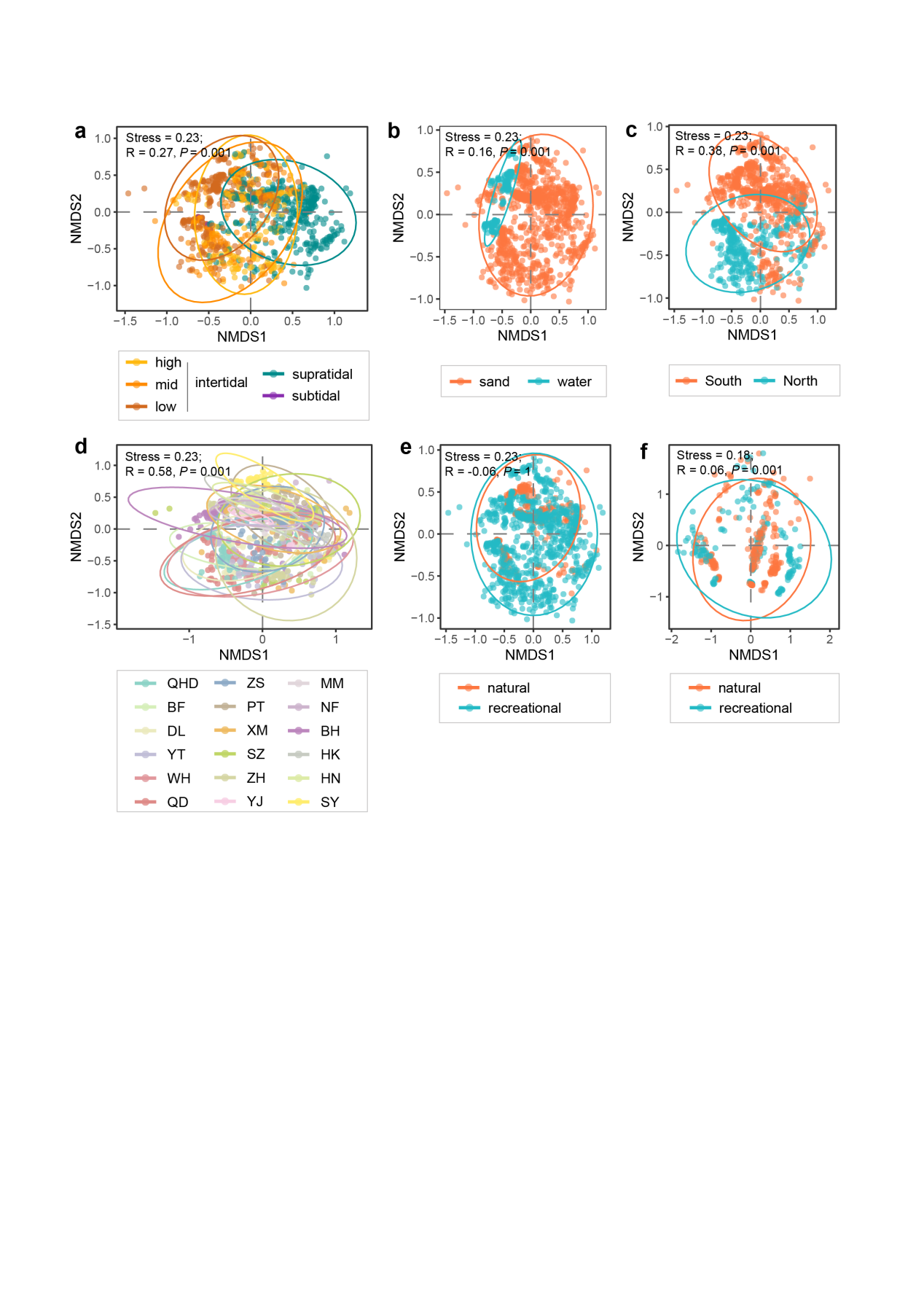

**Figure S5. Viral β-diversity across sandy beaches.** Non-metric multidimensional scaling (NMDS) ordinations based on Bray–Curtis dissimilarities showing viral community composition grouped by **(a)** tidal zones, **(b)** matrix type (sand versus water), **(c)** geographic regions (south versus north), **(d)** individual beaches (18 sites), and **(e–f)** beach usage (natural versus recreational). Panel **(e)** includes all samples, whereas panel **(f)** includes only paired recreational and adjacent natural beaches. Group differences were evaluated using ANOSIM with 999 permutations. Stress values, R statistics, and *P* values are shown in each panel.

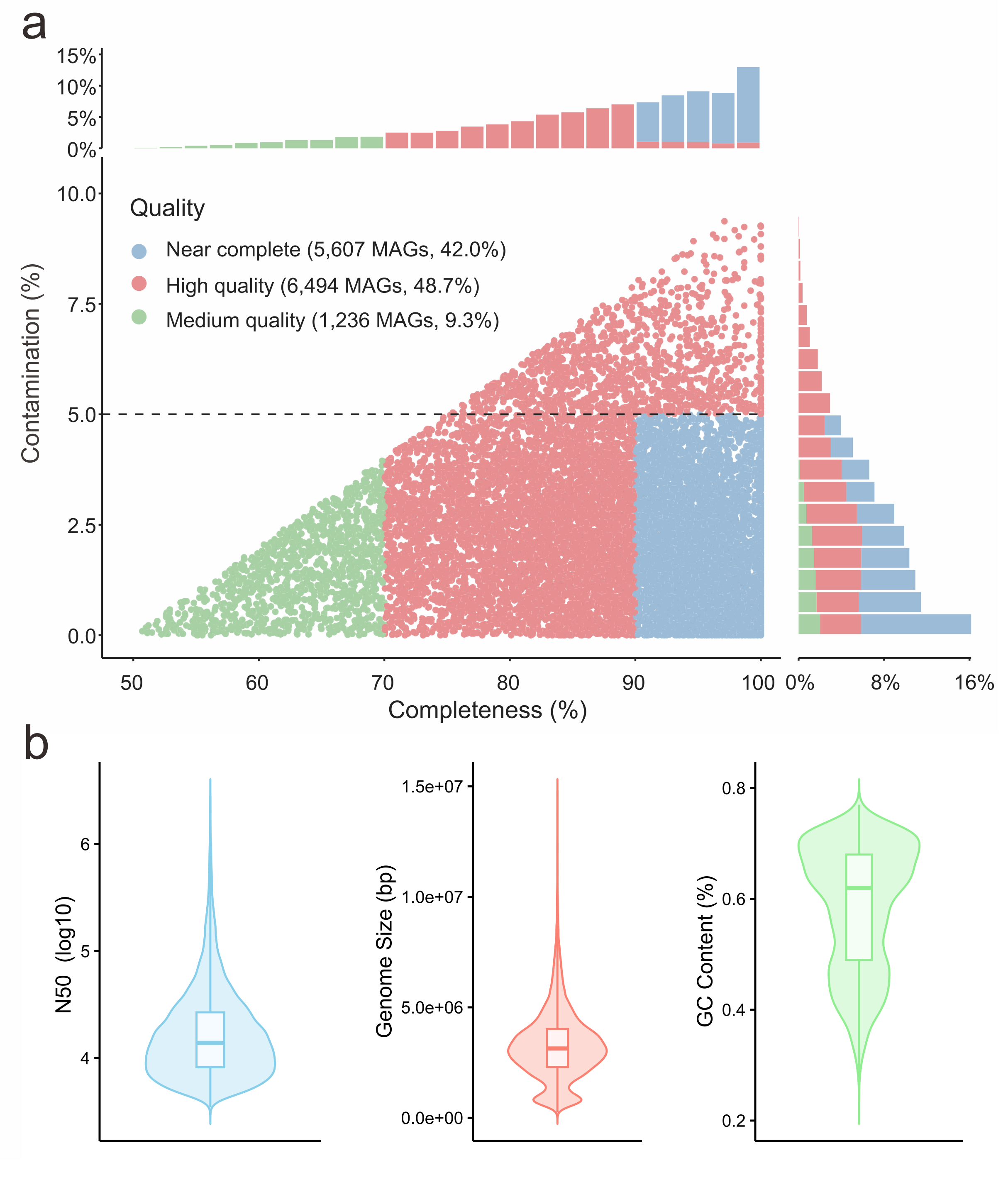

**Figure S6. Quality metrics of the Sandbeach Microbiome Genome Catalogue (SMGC). (a)** Distribution of genome completeness and contamination for 13,337 medium- to high-quality metagenome-assembled genomes (MAGs) comprising the SMGC. Points represent individual MAGs. Marginal bar plots show the percentage of MAGs across completeness (top) and contamination (right) intervals. Colors indicate quality categories (medium, near-complete, and high-quality). **(b)** Distributions of assembly statistics for all MAGs, including N50, genome size, and GC content. Boxes show the interquartile range with medians, and whiskers indicate 1.5× interquartile ranges. Detailed statistics are provided in **Table S16**.

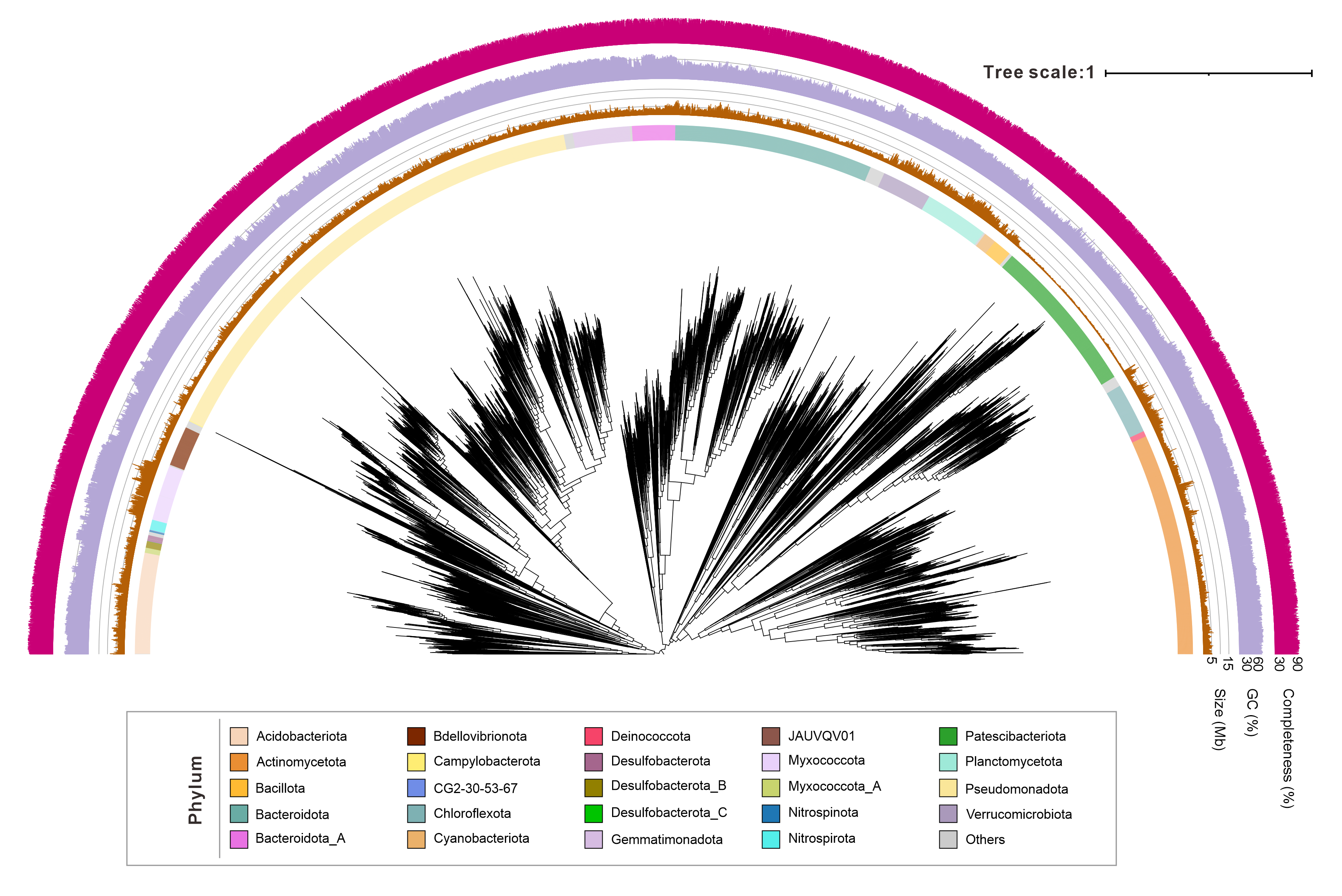

**Figure S7. Phylogenomic tree of bacterial metagenome-assembled genomes (MAGs) in the Sandbeach Microbiome Genome Catalogue (SMGC).** Maximum-likelihood tree reconstructed from 120 single-copy marker genes. Clades are colored by bacterial phylum. Concentric tracks from inner to outer indicate genome size, GC content, and completeness for each MAG.

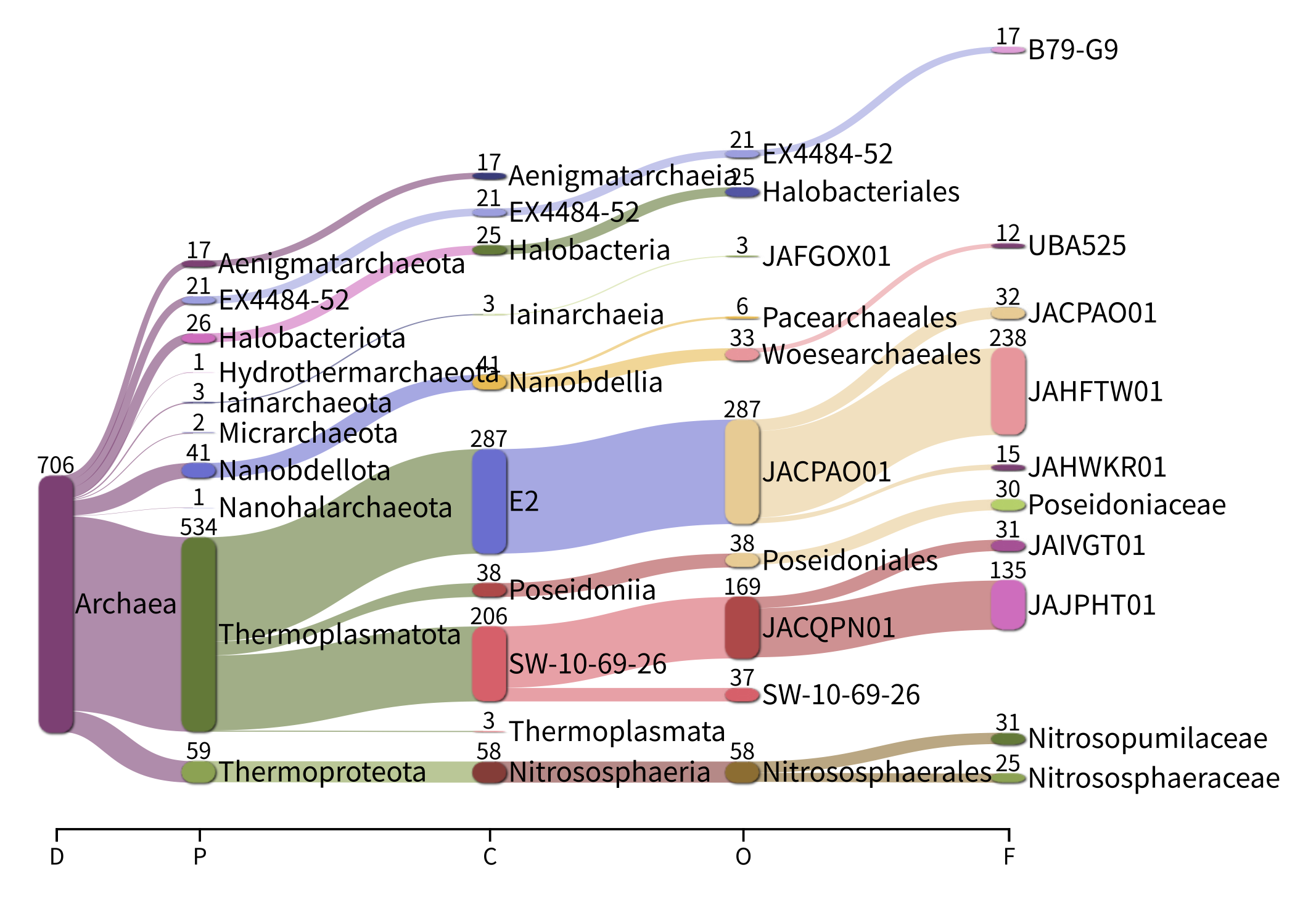

**Figure S8. Taxonomic composition of archaeal metagenome-assembled genomes (MAGs) in the Sandbeach Microbiome Genome Catalogue (SMGC).** Sankey diagram illustrating the distribution of archaeal MAGs across taxonomic ranks from domain to family based on GTDB r226 (D, domain; P, phylum; C, class; O, order; F, family). Flows represent the number of MAGs assigned to each lineage, and labels indicate counts.

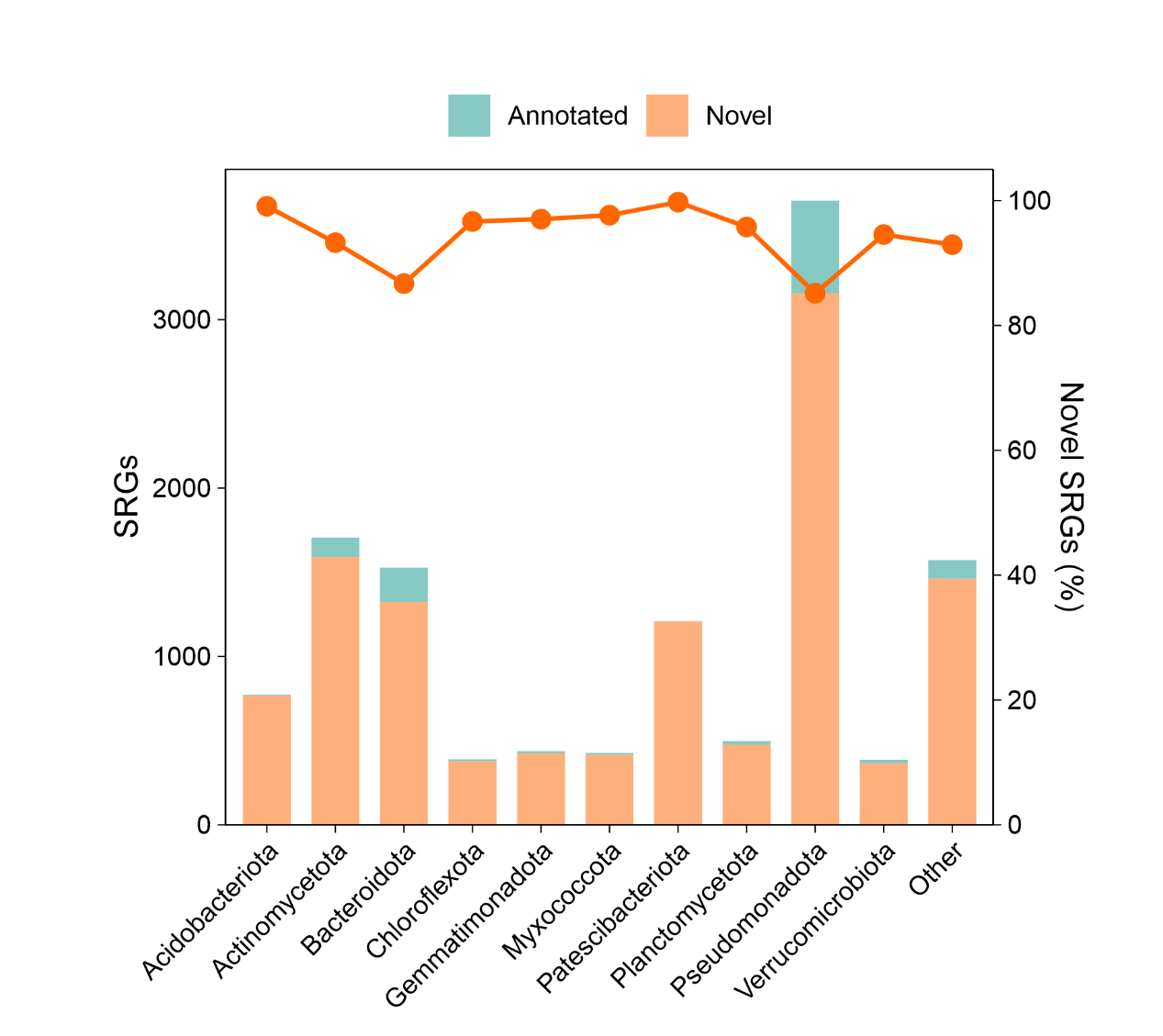

**Figure S9. Genomic novelty across dominant bacterial phyla in the Sandbeach Microbiome Genome Catalogue (SMGC).** Stacked bar plots show the number of species-representative genomes (SRGs) classified as annotated (previously described) or novel within each of the ten most abundant phyla. The line indicates the proportion of novel SRGs (% novelty) per phylum.

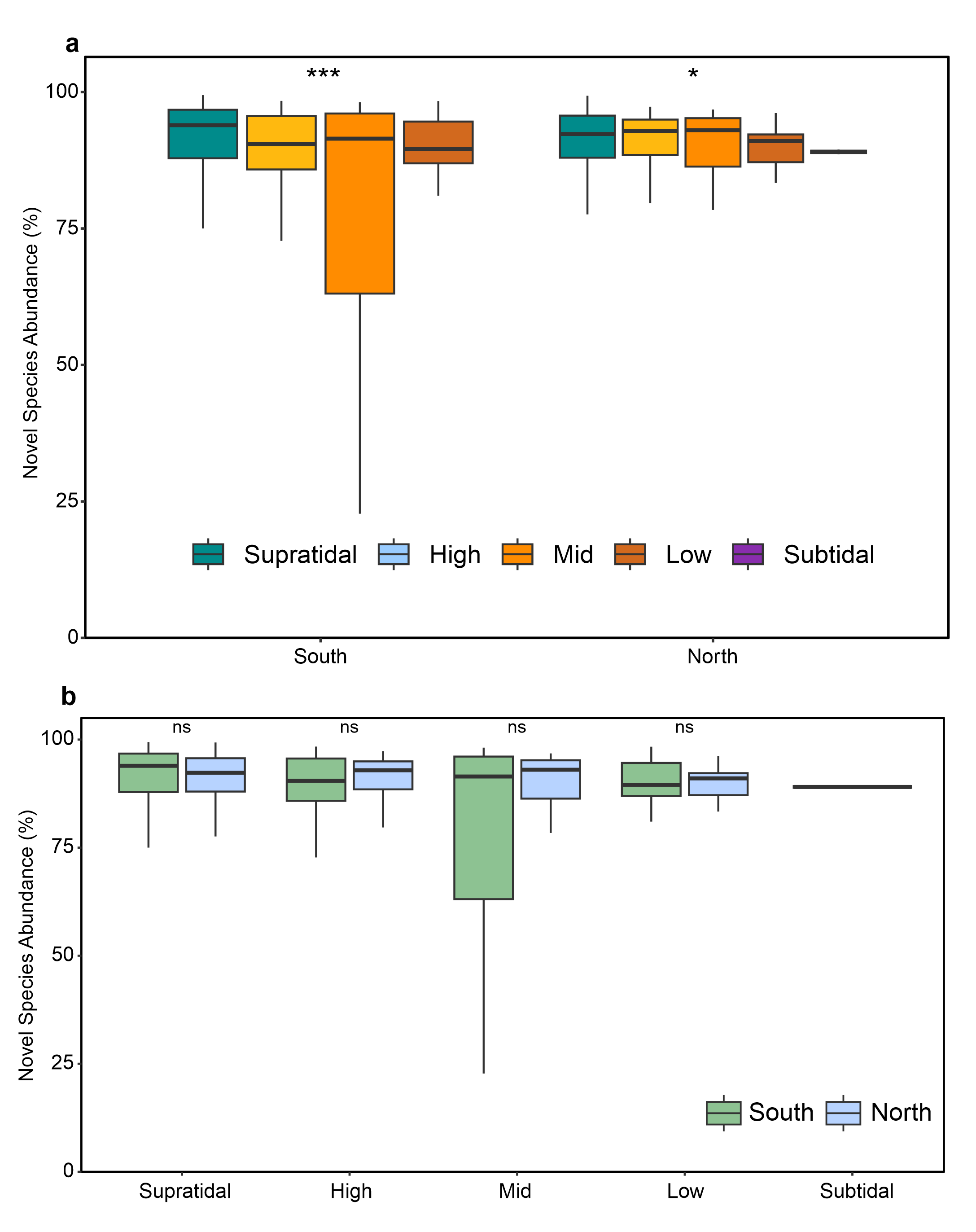

**Figure S10. Relative abundance of novel species-representative genomes (SRGs) across tidal zones and geographic regions. (a)** Novel SRG abundance compared among tidal zones in southern and northern beaches. **(b)** Novel SRG abundance compared between regions within each tidal zone. Boxplots show medians (center lines), interquartile ranges (boxes), and 1.5 × interquartile-range whiskers. Statistical differences were assessed using Kruskal–Wallis tests for tidal-zone comparisons and two-sided Wilcoxon rank-sum tests for regional comparisons. Significance is indicated above the boxes.

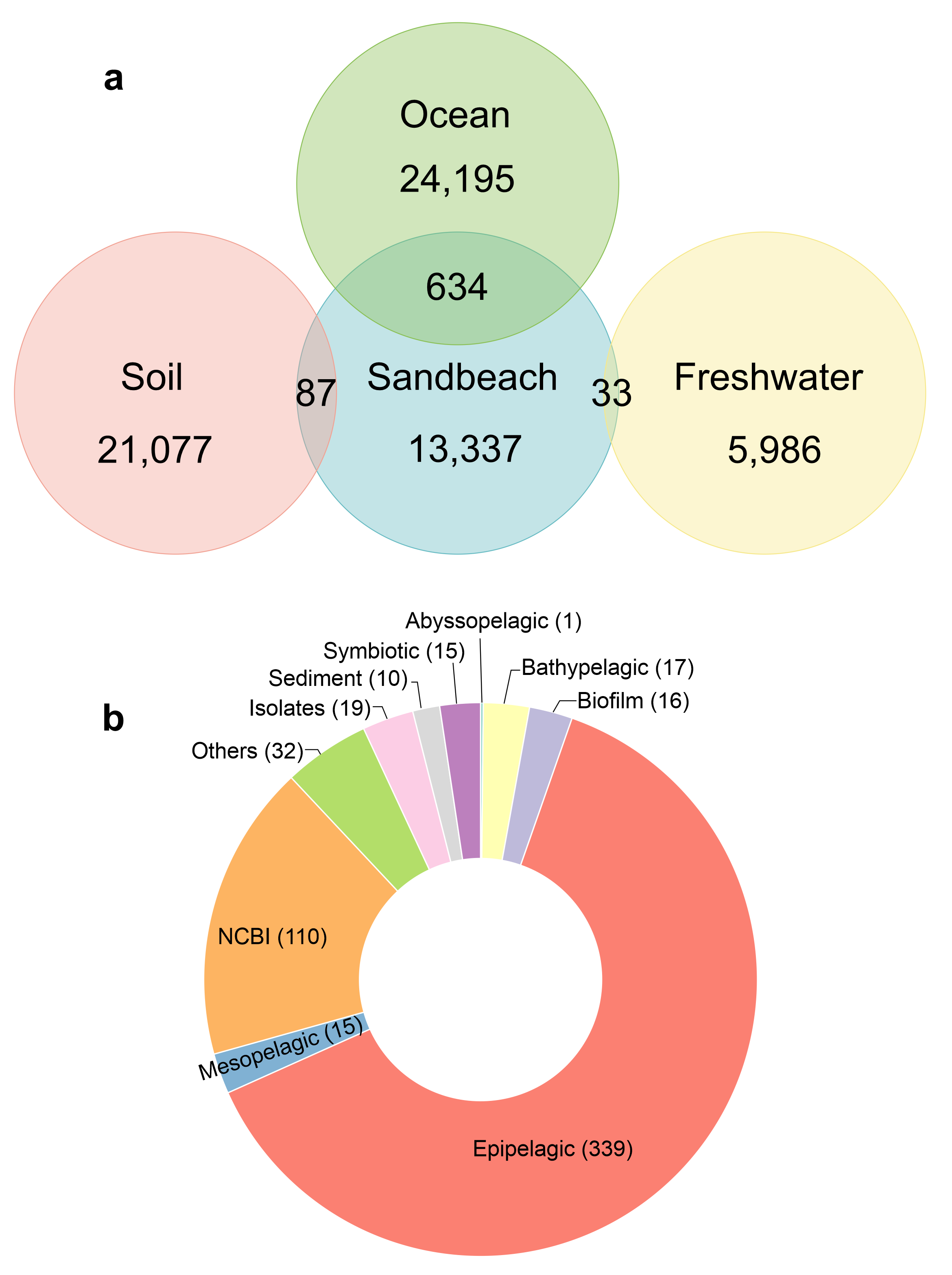

**Figure S11. Cross-habitat comparison of the Sandbeach Microbiome Genome Catalogue (SMGC). (a)** Venn diagram showing overlap of species-representative genomes (SRGs) between the SMGC and genome collections from ocean, soil, and freshwater habitats. Numbers indicate shared and habitat-specific genomes. **(b)** Donut chart showing the environmental origins of ocean–sandy beach shared SRGs, with segment sizes representing counts per habitat category.

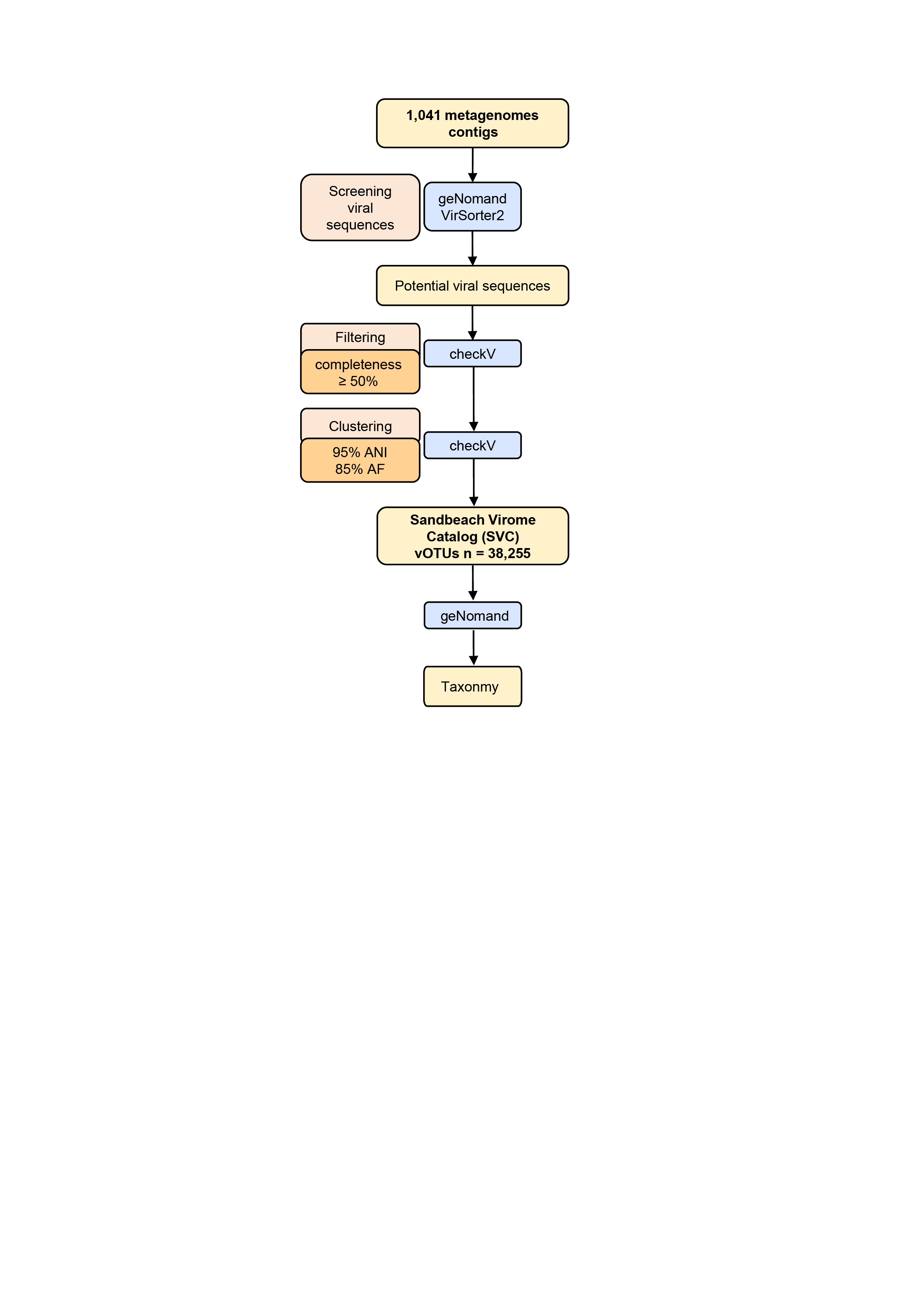

**Figure S12. Workflow for construction of the Sandbeach Virome Catalogue (SVC).** Metagenomic contigs were screened for viral sequences using geNomad and VirSorter2. Candidate viral contigs were quality-filtered (≥50% completeness; CheckV) and clustered into viral operational taxonomic units (vOTUs) at 95% average nucleotide identity and 85% alignment fraction. The resulting 38,255 vOTUs were taxonomically annotated using geNomad.

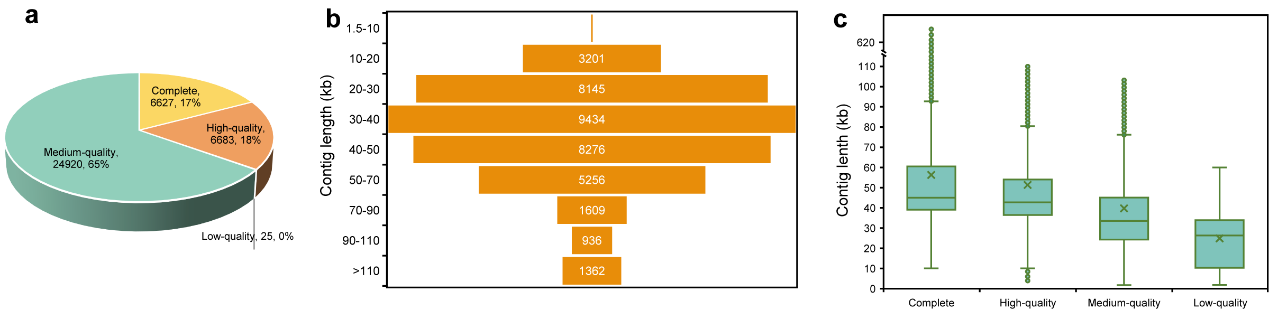

**Figure S13. Quality assessment of viral operational taxonomic units (vOTUs) in the Sandbeach Virome Catalogue (SVC). (a)** Proportion of vOTUs across completeness categories as estimated by CheckV. **(b)** Distribution of vOTU lengths. **(c)** Length distributions stratified by completeness category. Detailed information for individual vOTUs is provided in **Table S18**.

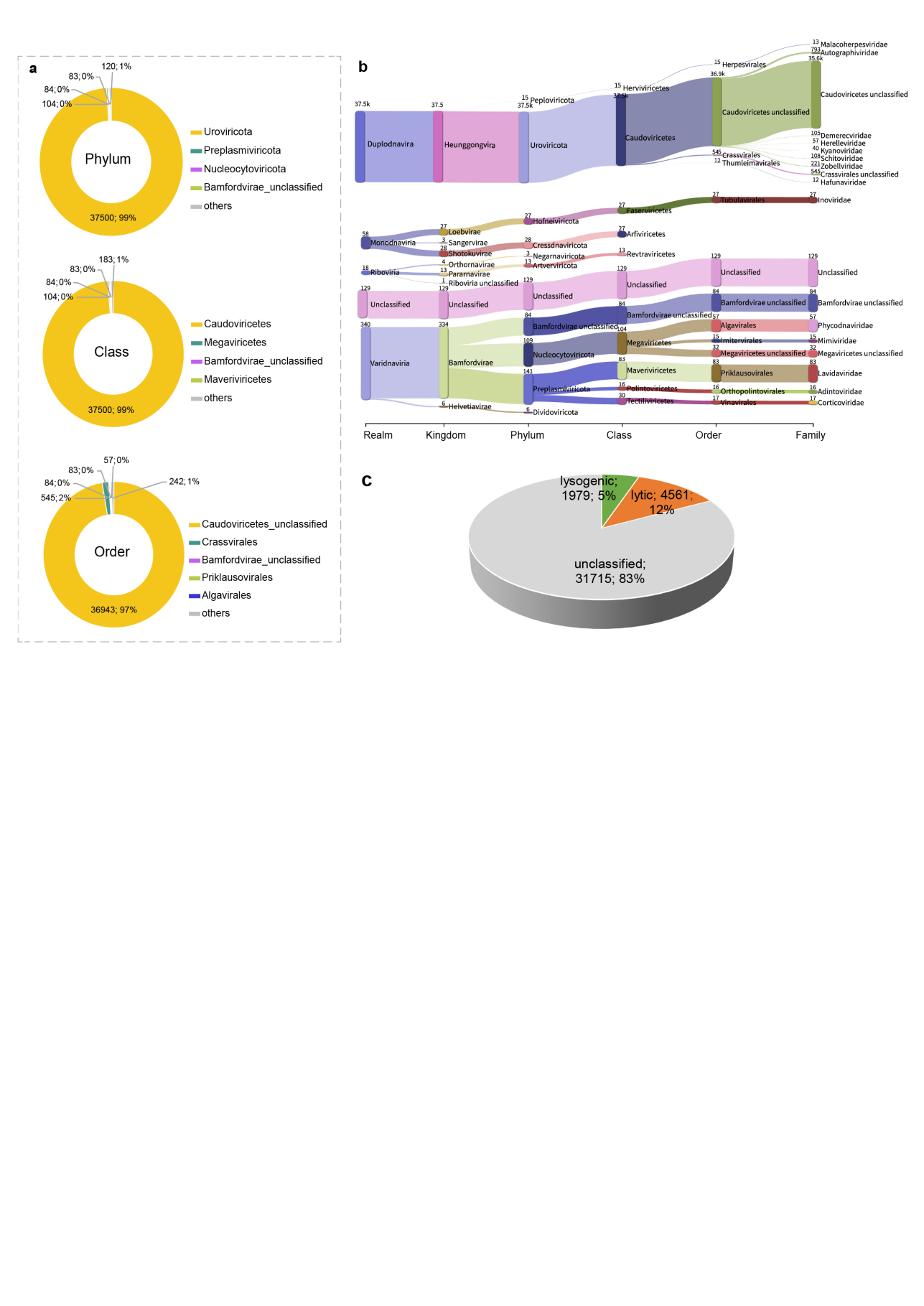

**Figure S14. Taxonomic composition and predicted lifestyles of vOTUs in the Sandbeach Virome Catalogue (SVC). (a)** Proportional distributions of classified vOTUs at the phylum, class, and order levels. **(b)** Sankey diagram showing taxonomic assignments from realm to family, with flow widths representing the number of vOTUs at each rank. **(c)** Predicted viral lifestyles (lytic, lysogenic, and unclassified). Detailed taxonomic and lifestyle statistics are provided in **Tables S19** and **S22**.

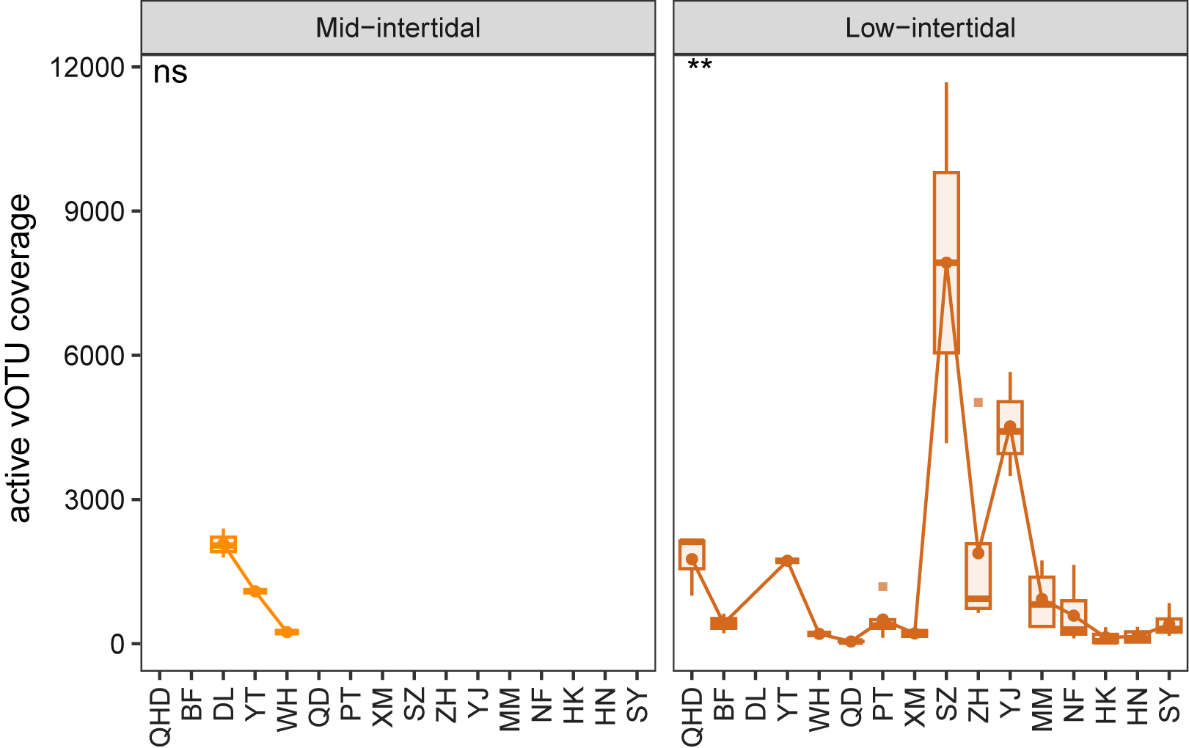

**Figure S15. Coverage of transcriptionally active vOTUs in the Sandbeach Virome Catalogue (SVC).** Boxplots show transcript-derived coverage of active vOTUs in water samples from the mid-intertidal (left) and low-intertidal (right) zones, ordered latitudinally from north to south. Significance between regions is indicated above the panels. Detailed values are provided in **Table S20**.

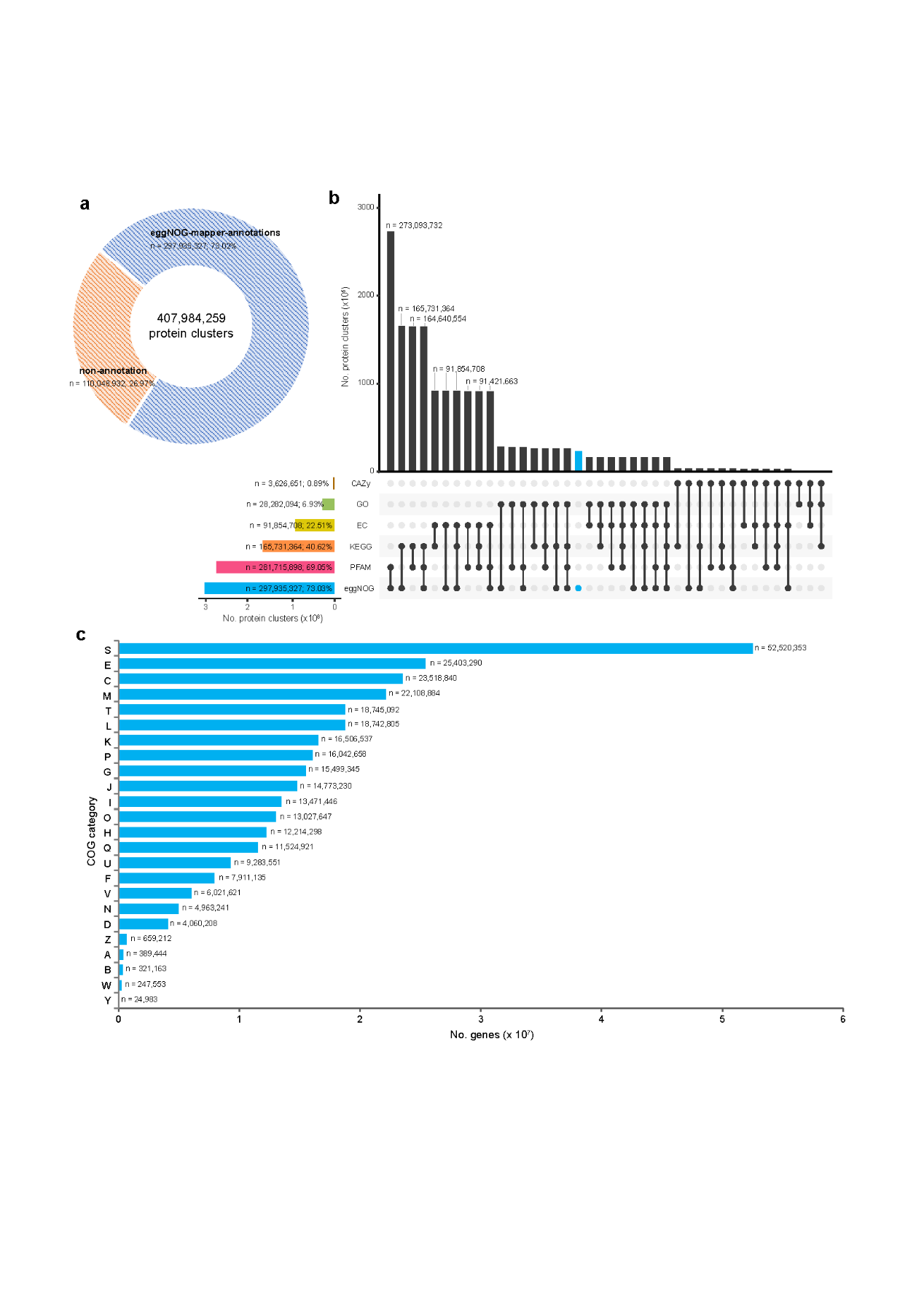

**Figure S16. Functional annotation landscape of the sandy beach non-redundant gene catalogue. (a)** Proportion of protein clusters with functional annotations. **(b)** Overlap of annotations across databases (eggNOG, Pfam, KEGG, EC, GO, and CAZy); bars indicate the numbers of protein clusters annotated by individual databases or their combinations. **(c)** Distribution of annotated genes across Clusters of Orthologous Groups (COG) functional categories.

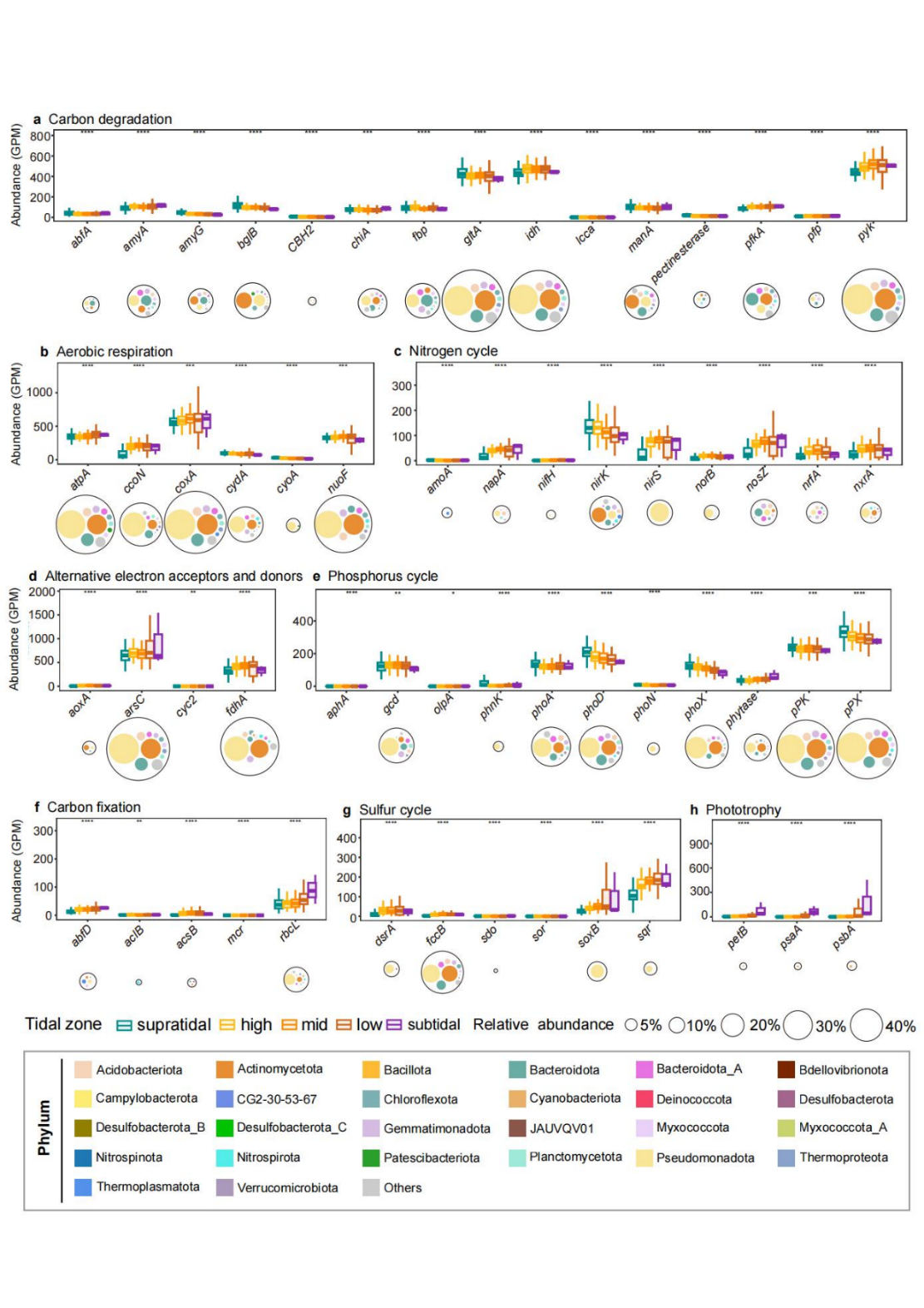

**Figure S17.** Metabolic potential of sandy beach microbial communities across tidal zones. Boxplots show the abundance of functional marker genes (genes per million, GPM) involved in **(a)** carbon degradation, **(b)** aerobic respiration, **(c)** nitrogen cycling, **(d)** alternative electron acceptors and donors, **(e)** phosphorus cycling, **(f)** carbon fixation, **(g)** sulfur cycling, and **(h)** phototrophy. Boxplots show medians (center lines), interquartile ranges (boxes), and 1.5 × interquartile-range whiskers. Colors denote tidal zones. Statistical differences were evaluated using Kruskal–Wallis tests, with significance indicated as ns, **P* < 0.05, ***P* < 0.01, and ****P* < 0.001. Circles represent MAGs encoding the corresponding pathways; circle size indicates average relative abundance and color denotes phylum.

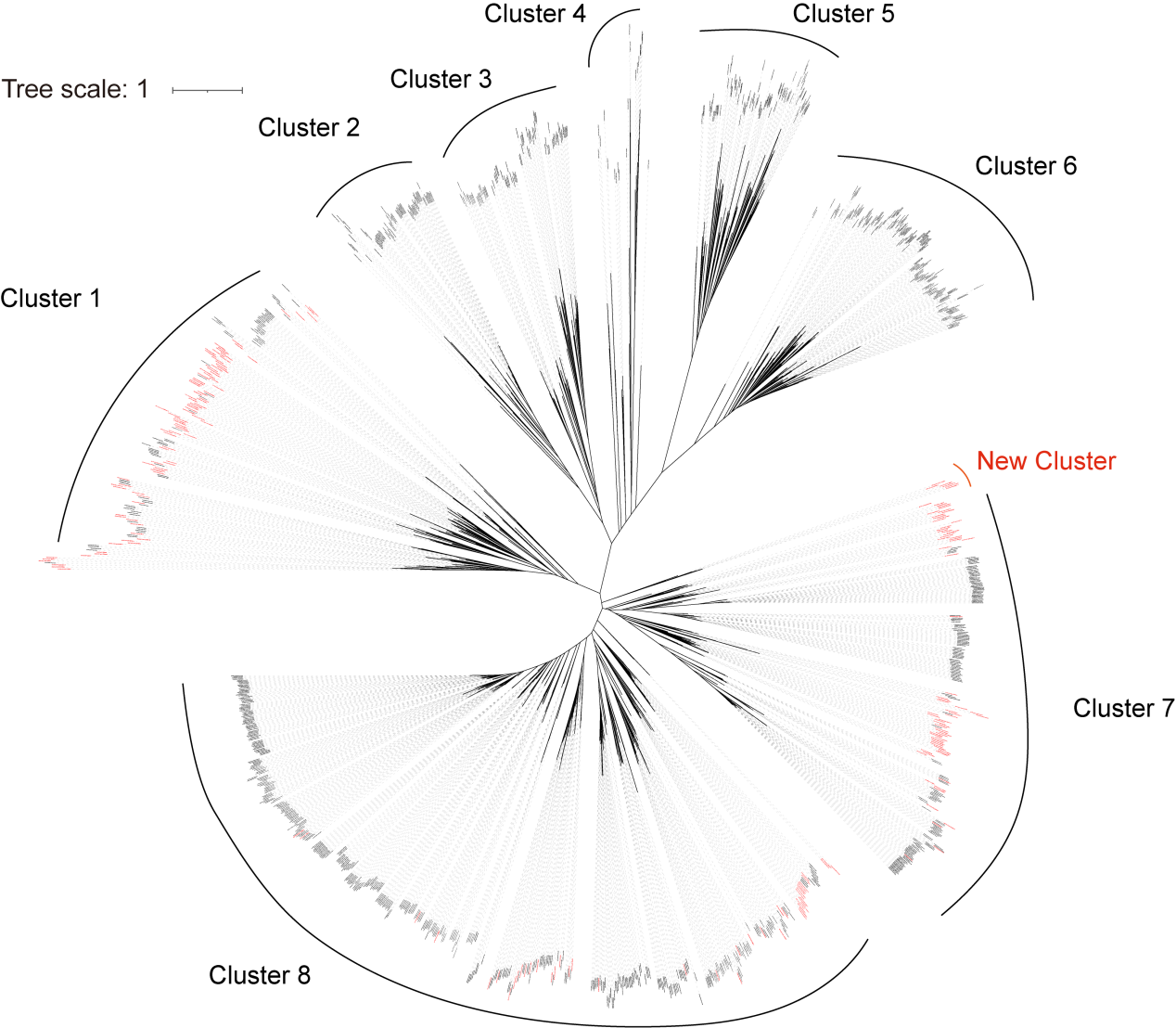

**Figure S18. Phylogeny of alkane monooxygenases (AlkB).** Maximum-likelihood tree inferred from amino acid sequences. Branch lengths indicate substitutions per site. Sequences recovered in this study are highlighted in red, whereas reference sequences are shown in black. Major clades are indicated, including a newly identified cluster.

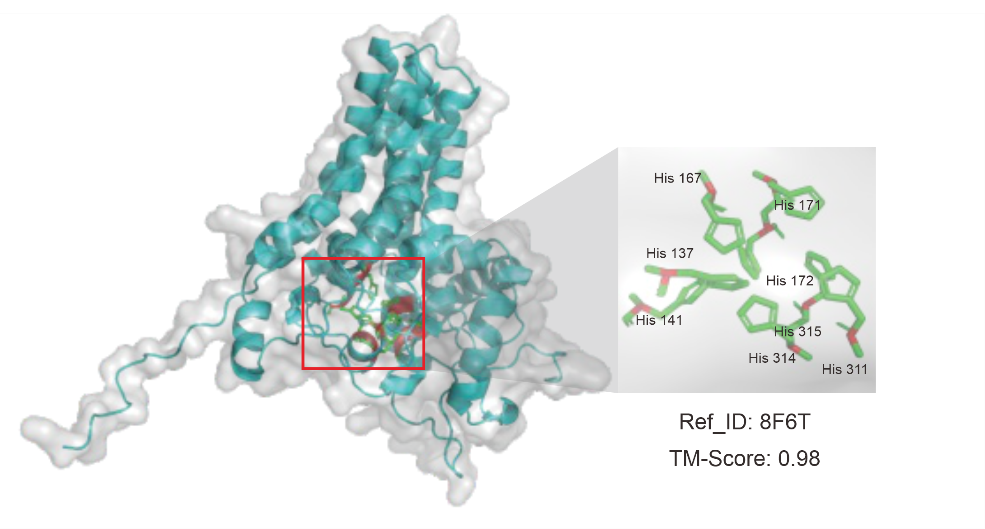

**Figure S19. Structural alignment of predicted AlkB with an experimentally validated AlkB structure (PDB: 8F6T, gray).** Catalytic residues are shown as sticks.

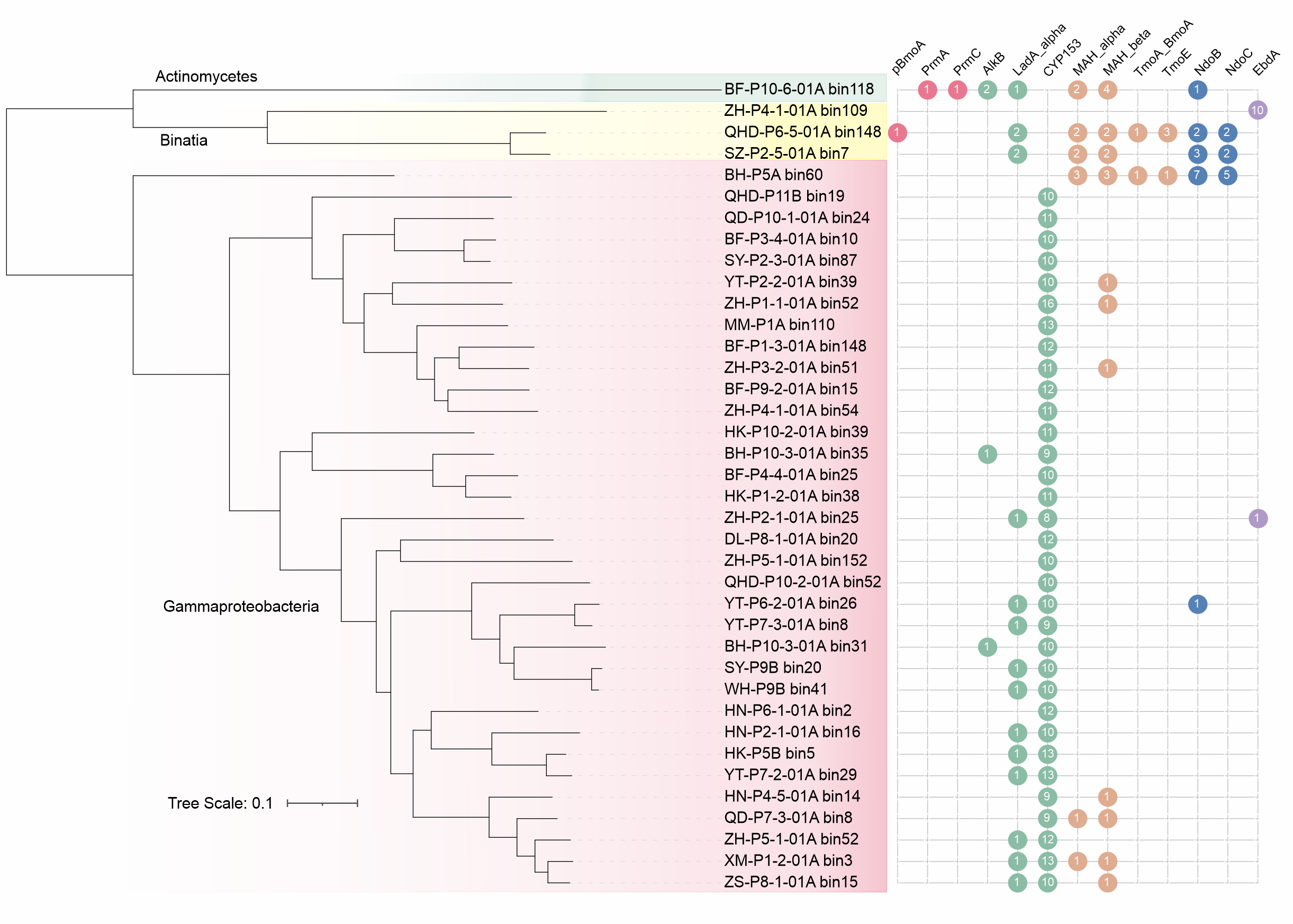

**Figure S20. Phylogenetic distribution of hydrocarbon-degrading potential across sandy beach genomes.** Maximum-likelihood genome tree reconstructed from concatenated alignments of 120 single-copy marker genes for representative hydrocarbon-degrading bacteria. Branch lengths indicate substitutions per site. The matrix shows the presence and copy number of key hydrocarbon degradation genes across genomes, with circle size proportional to gene counts.

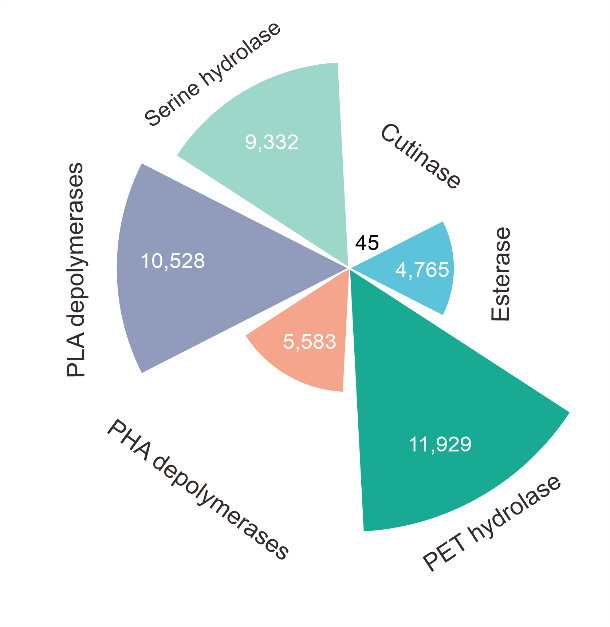

**Figure S21. Distribution of plastic-degrading genes in the Sandbeach Microbiome Genome Catalogue.** Numbers of plastic-degrading genes are shown in white. Detailed annotations are provided in **Table S27**.

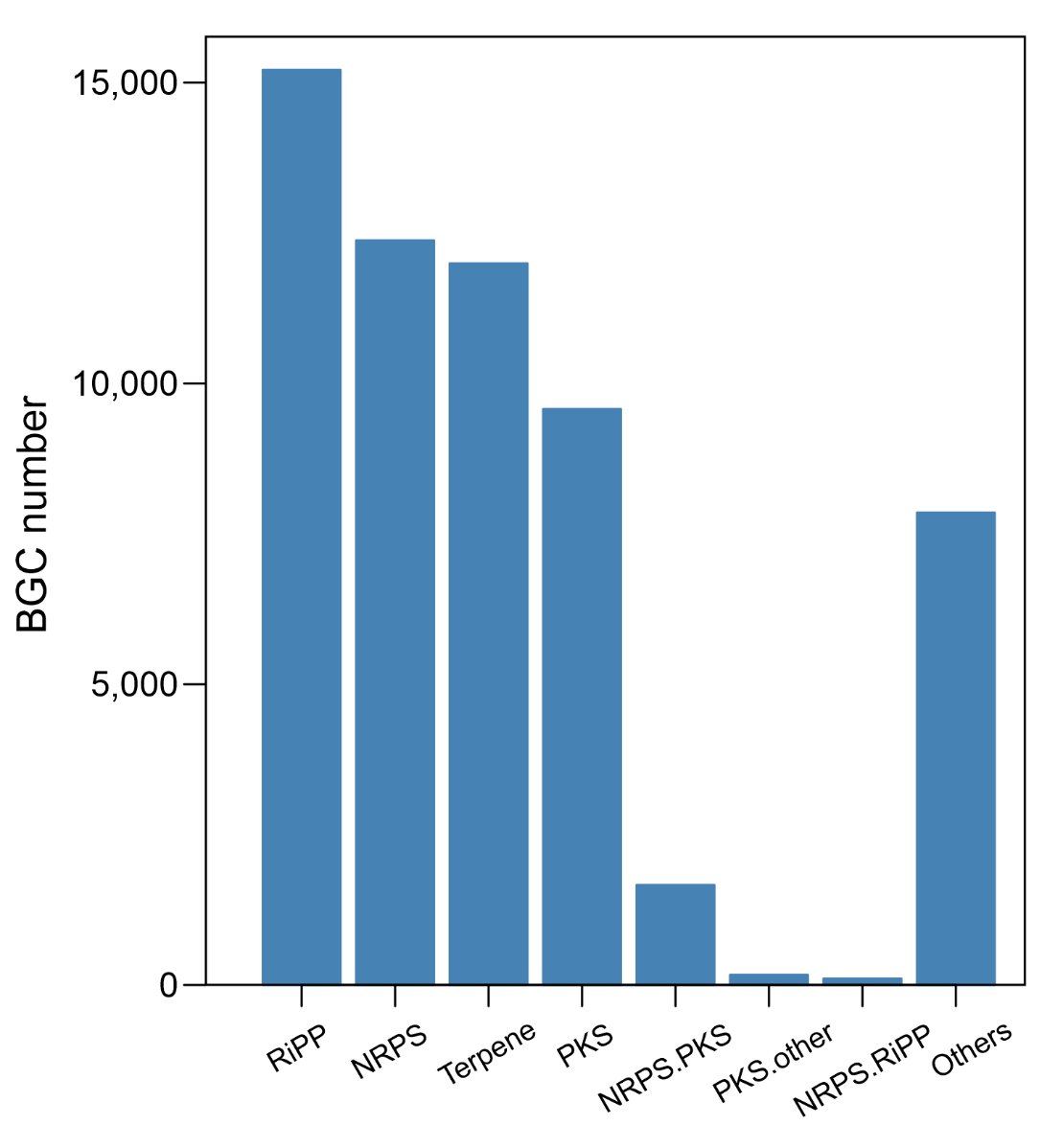

**Figure S22. Distribution of biosynthetic gene cluster (BGC) classes in the Sandbeach Microbiome Genome Catalogue (SMGC).** Bar plot showing the number of BGCs assigned to each biosynthetic class. Detailed annotations are provided in **Table S32**.

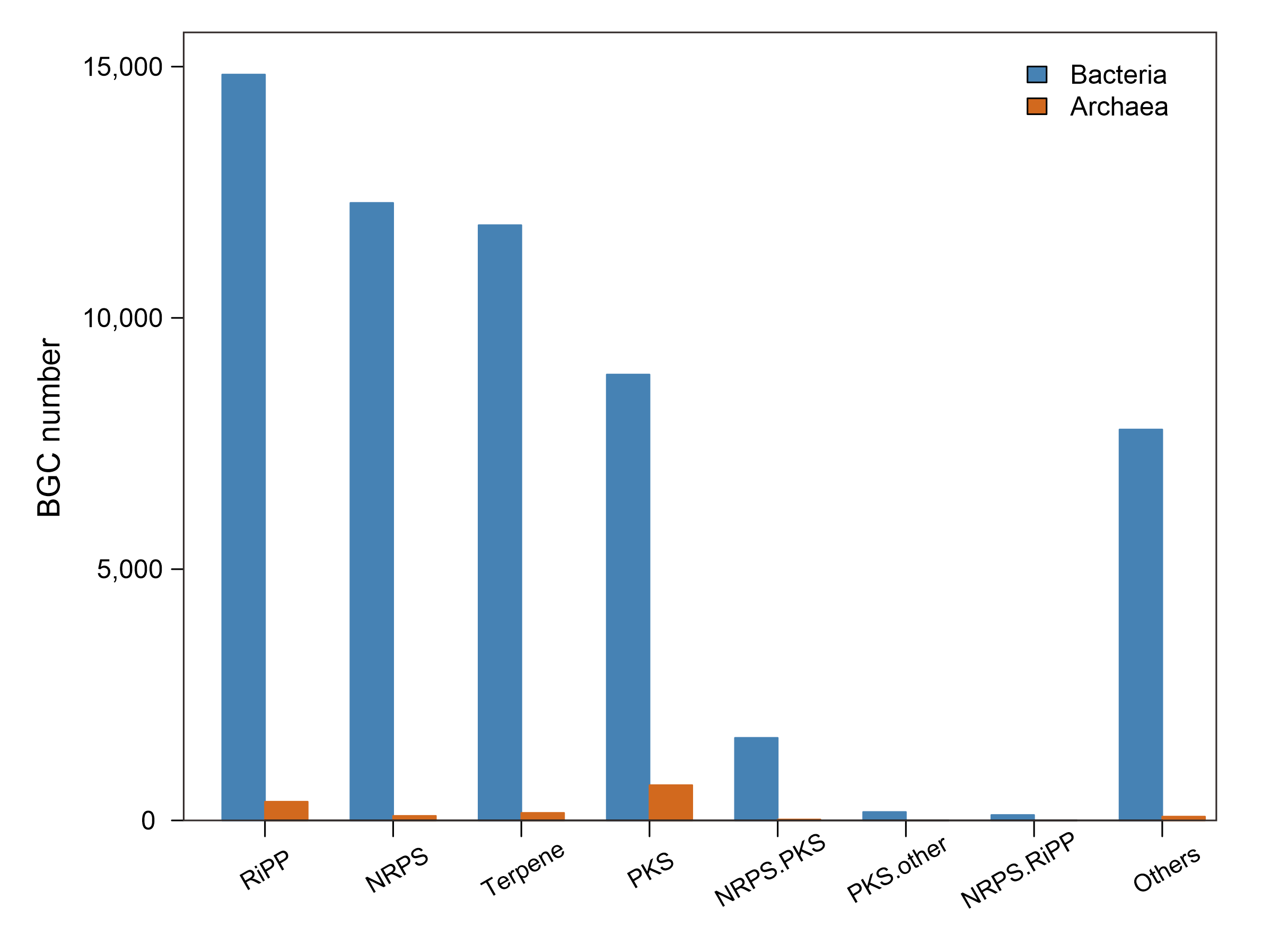

**Figure S23.** **Distribution of biosynthetic gene cluster (BGC) classes in bacterial and archaeal MAGs.** Bar plot showing the number of predicted BGCs assigned to each biosynthetic class for bacteria and archaea. Detailed annotations are provided in **Table S32**.
